# The conserved, secreted protease inhibitor MLT-11 is necessary for *C. elegans* molting and embryogenesis

**DOI:** 10.1101/2022.06.29.498124

**Authors:** James Matthew Ragle, Max T. Levenson, John C. Clancy, An A. Vo, Vivian Pham, Jordan D. Ward

**Affiliations:** Department of Molecular, Cell, and Developmental Biology, University of California-Santa Cruz, Santa Cruz, CA 95064, USA

**Keywords:** *C. elegans*, molting, Kunitz domain, apical extracellular matrix, protease inhibitor

## Abstract

Apical extracellular matrices (aECMs) are associated with all epithelia and form a protective layer against biotic and abiotic threats in the environment. *C. elegans* molting offers a powerful entry point to understanding developmentally programmed aECM remodeling. Several protease inhibitors are implicated in molting, but their functions remain poorly understood. Here we characterize *mlt-11*, an unusual protease inhibitor with 10 conserved Kunitz domains. MLT-11 oscillates and is localized in the cuticle and in lysosomes in larvae and in the embryonic sheath starting at the 3-fold embryo stage. *mlt-11* (RNAi) produced a developmental delay, motility defects, failed apolysis, and a defective cuticle barrier. *mlt-11* null and C-terminal Kunitz domain deletion mutants are embryonic lethal while N-terminal deletions cause a rolling phenotype indicative of cuticle structure abnormalities. *mlt-11* activity is primarily necessary in seam and hypodermal cells and accordingly *mlt-11* (RNAi) causes defects in localization of the collagens ROL-6 and BLI-1 over the cuticle. *mlt-11* (RNAi) molting phenotypes can be suppressed by genetically inhibiting endocytosis. Our model is that MLT-11 is acting in the aECM to coordinate remodeling and timely ecdysis.

## INTRODUCTION

Specialized extracellular matrices cover the apical surface of all epithelial cells and form the skin in almost all animals (Li Zheng et al., 2020). These apical extracellular matrices (aECMs) also line the lumen of internal tubular epithelia to form a protective layer against biotic and abiotic threats (Li Zheng et al., 2020). Despite their importance, understanding the structure and dynamics of aECM components in development and disease remains challenging.

*C. elegans* is emerging as a powerful model to study aECM structure and remodeling. They have a collagen-based ECM so understanding its assembly may provide insight into mammalian skin (Page and Johnstone, 2007). The components of the cuticle are secreted by hypodermal and seam cells and are assembled in distinct layers (Page and Johnstone, 2007). During each larval stage animals must build a new aECM underneath the old one, separate the old aECM (apolysis) and then shed it (ecdysis) (Laźetić and Fay, 2017b). A specialized, transient structure known as the pre-cuticle is thought to pattern the new cuticle and is then shed during ecdysis (Cohen and Sundaram, 2020). The sheath is a similar structure in embryos which ensures embryonic integrity and directs force during elongation (Vuong-Brender et al., 2017). The vulval aECM has recently been shown to be highly dynamic, and specialized aECMs also line the rectum and excretory system (Cohen et al., 2019; Cohen et al., 2020; Gill et al., 2016).

A major question is how is the aECM remodeled during molting? Proteases are required for ecdysis in both *C. elegans* and in parasitic nematodes, presumably by promoting apolysis, though some are thought to function in collagen processing (Davis et al., 2004; Hashmi et al., 2004; Kim et al., 2011; Stepek et al., 2011). Protease inhibitors have been implicated in molting through RNAi screening, and have been suggested to suppress ecdysis (Frand et al., 2005; Laźetić and Fay, 2017b). BLI-5 has homology to the Kunitz domain family of protease inhibitors and mutations cause molting defects. However, recombinant BLI-5 enhanced the activity of two serine proteases from distinct classes (Page et al., 2006; Stepek et al., 2010). MLT-11 is another putative protease inhibitor in the Kunitz family and *mlt-11 (RNAi)* causes molting defects (Frand et al., 2005). *mlt-11* mRNA oscillates, peaking mid molt and its expression is regulated by NHR-23, a nuclear hormone receptor transcription factor necessary for molting (Frand et al., 2005). However, *mlt-11* remains poorly characterized.

Here we demonstrate that MLT-11 is localized to the cuticle, lysosomes, the rectal epithelium, excretory duct lumen, and vulval lumen. It is also secreted into the extracellular space in embryos before localizing to the cuticle prior to hatching. *mlt-11* null alleles cause embryonic lethality, characterized by disorganization of adherens junctions. RNAi in larvae causes developmental delay, apolysis, and ecdysis defects. *mlt-11* activity in seam cells is necessary for molting and a normal developmental rate. *mlt-11* inactivation causes a defective cuticle barrier and aberrant localization of the collagens ROL-6 and BLI-1. Genetic data suggests that MLT-11 acts in the aECM. This work provides the first insight into how MLT-11 functions to promote embryogenesis and aECM integrity during molting.

## RESULTS

### *mlt-11* is expressed in embryonic, larval, and adult epidermal cells

*mlt-11* has been reported to be an NHR-23 target gene (Frand et al., 2005). There are four NHR-23 ChIP-seq peaks in the *mlt-11* promoter (Johnson et al., 2022), and the sequences under these peaks are highly conserved in other nematodes (Fig. 1A; see Conservation track). There are additional areas of the promoter which display elevated conserved sequence, which may indicate other regulatory elements (Fig. 1A). We used 2.8 kilobases of upstream sequence to create a single copy *mlt-11p::NLS::mNeonGreen* promoter reporter. We note that this sequence differs in part from the promoter used by Frand et al., 2005 (Fig 1A). Expression in embryos was first detected at the bean stage in posterior epithelial cells and persisted through the 3-fold stage spreading more anteriorly (Fig. 1C). Expression was detected in hypodermal, rectal and vulval cells in both larvae and adults as well as seam cells in larvae (Fig. 1B).

**Fig. 1.**
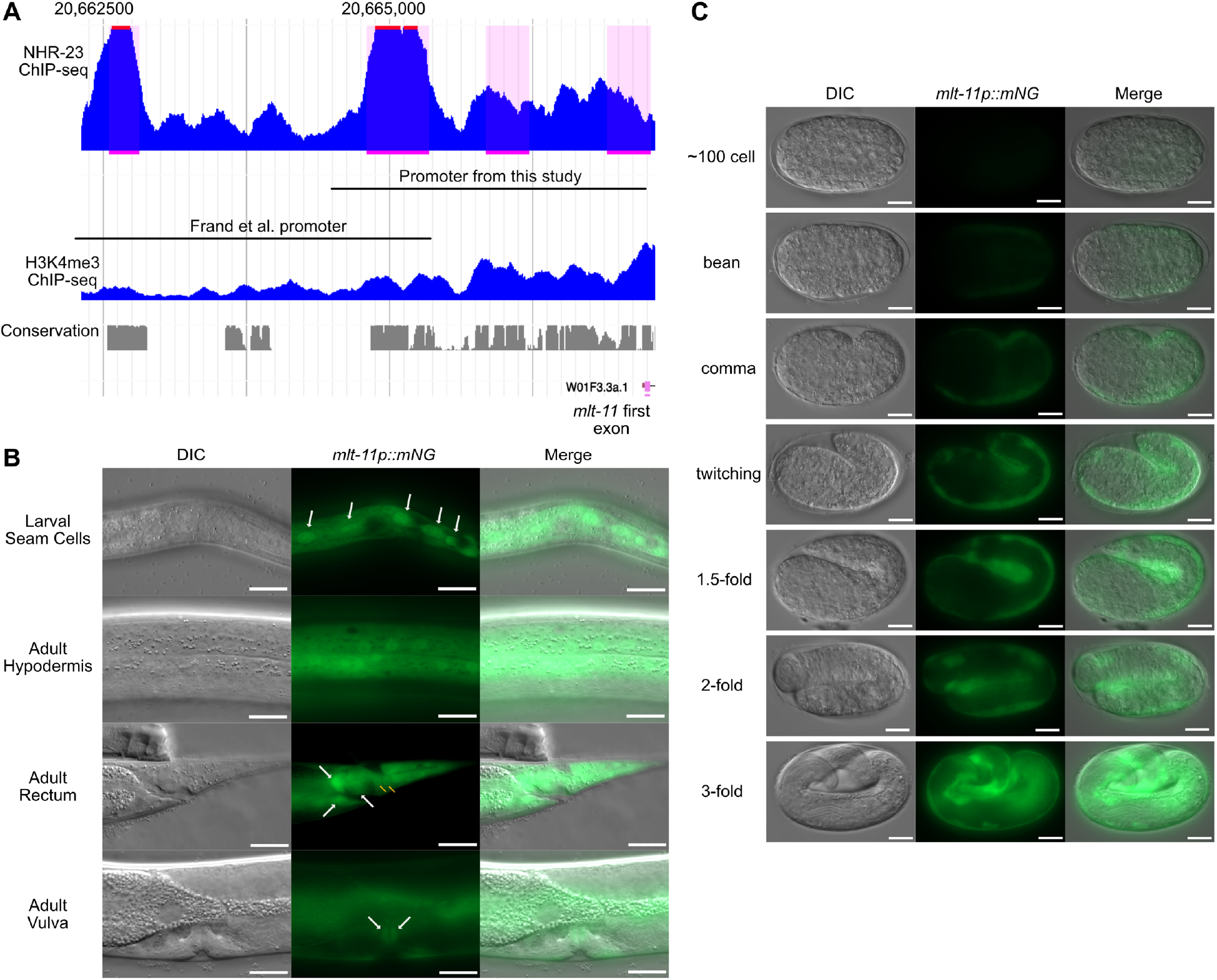
*mlt-11* is expressed in larval and embryonic epidermal cells. (A) Genome browser track of *mlt-11* promoter depicting NHR-23 and H3K4me3 ChIP-seq peaks and conservation calculated across 26 nematode species. The promoter used in Fig. 1B and 1C and Frand et al. (2005) are indicated. Genomic sequence on chromosome V is indicated above. *mlt-11p::mNeonGreen (mNG)*, DIC, and overlay images of the indicated larval and adult (B) and embryonic (C) stages. In B, white arrows indicate seam cells (top), rectal epithelial cells (middle) and vulval cells (bottom) and yellow arrows indicate hypodermal cells near rectum. Images are representative of 40 animals examined over two biological replicates. Scale bars in B are 20 µm and in C are 10 µm.

### MLT-11 is an oscillating secreted protein that localizes to the aECM and lysosomes

To determine where MLT-11 localized, we knocked an *mNeonGreen::3xFLAG* cassette into the 3’ end of the gene producing a C-terminal translational fusion that labels all described *mlt-11* isoforms (Fig. 2A). MLT-11::mNeonGreen::3xFLAG (MLT-11::mNG) was detected in the excretory duct, hypodermal cells, seam cells and the rectum (Fig. 2B). In the hypodermis MLT-11::mNG was non-nuclear and either diffuse through the cytoplasm or in bright punctae, lysosomal based on morphology (Miao et al., 2020). The cytoplasmic expression was reminiscent of secreted proteins. We confirmed that this pattern reflected localization to the endoplasmic reticulum through co-localization with an mCherry::TRAM marker (Fig. 2C)(Chen et al., 2012). MLT-11::mNG localization in the vulva was dynamic. In early L4 (stage 4.3 by vulva morphology) MLT-11::mNG was lumenal and by mid-L4 (stage 4.5) we saw expression within the vulD cell (Fig. 2D). In late L4 (stage 4.8) MLT-11::mNG was robustly expressed in vulD and in the vulval lumen (Fig. 2D). In embryos, MLT-11::mNG was first observed at the bean stage (Fig. 2E). From this stage to the 3-fold stage MLT-11::mNG appeared to be secreted, localizing in the space between the embryo and the eggshell with enrichment at the embryo epidermis (Fig. 2E). During elongation, MLT-11::mNG perinuclear expression was observed, reminiscent of the collagen DPY-7 localization to embryonic endoplasmic reticulum (McMahon et al., 2003). Before hatching there was a striking shift in MLT-11::mNG localization where it labeled annuli in the aECM (Fig. 2E). Immunoblotting revealed that MLT-11 oscillates, and three isoforms were detected (Fig. 2F). Two isoforms are large (260 kDa) and align with the predicted size of full-length MLT-11 isoforms with an mNG tag (268.5-371.4 kDa); these isoforms peak in early L4 and rapidly disappear (Fig. 2H). A ∼50-70 kDa band smaller than any predicted isoform appears in early L4 and persists until late L4 (Fig. 2H). Together, these data indicate that MLT-11 is an oscillating secreted protein with dynamic localization to aECM in embryos and larvae.

**Fig. 2.**
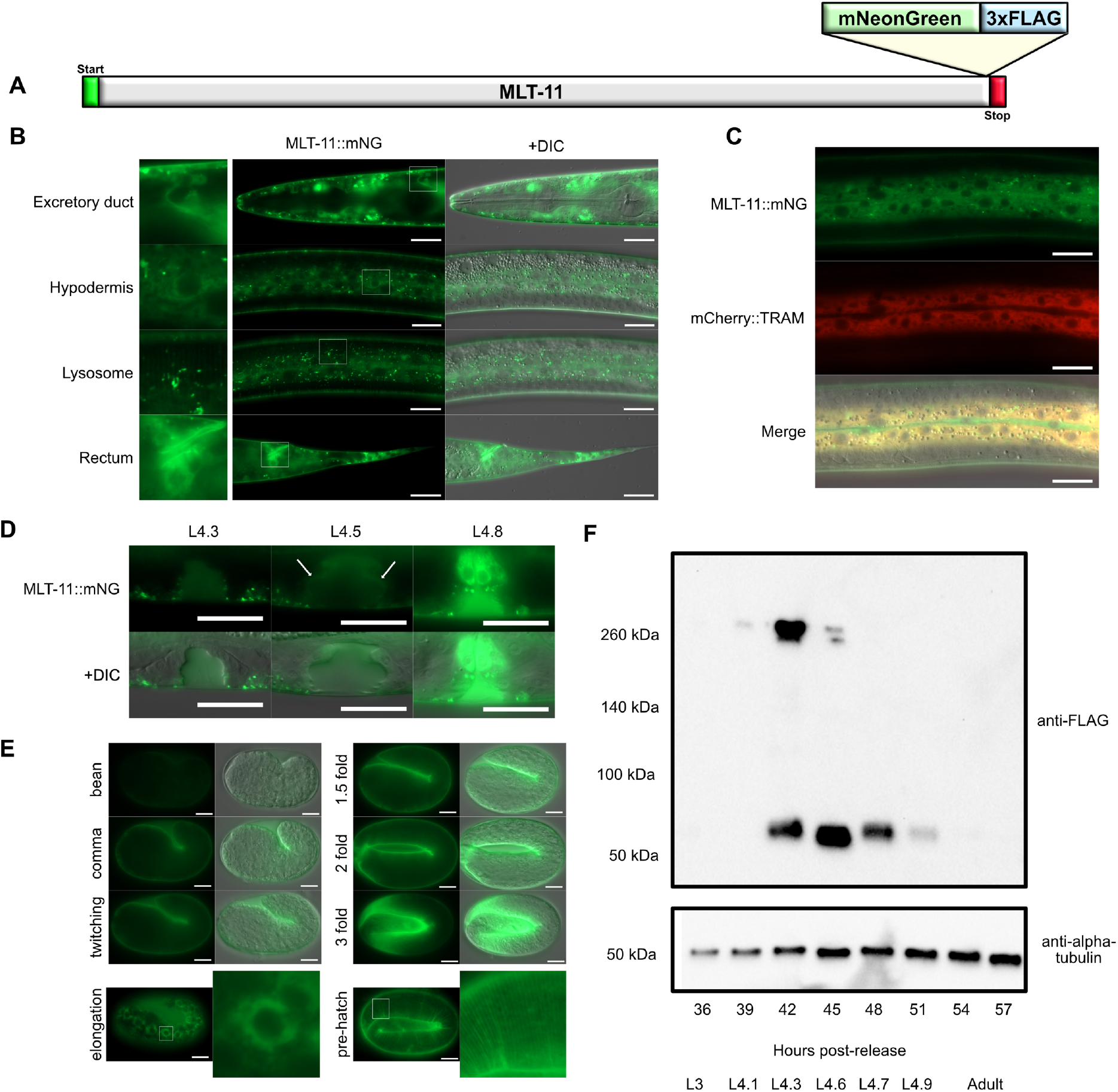
MLT-11 localization. (A) Schematic of *mlt-11::mNG::3xFLAG* knock-in. (B) MLT-11::mNG localization in the indicated tissues along with DIC overlay. Boxes on the left are magnified cellular or subcellular structures within the larger images to the right. (C) MLT-11::mNG overlay with mCherry::TRAM. (D) MLT-11::mNG vulval localization in the indicated L4 stages. (E) MLT-11::mNG localization in the indicated embryonic stages. Images are representative of 40 animals examined over two biological replicates. Scale bars in B and C are 20 µm and in E are 10 µm. (H) Immunoblotting with the indicated antibodies of *mlt-11::mNG::3xFLAG* lysates harvested at the indicated time points post-release. A developmental stage for each time point as determined by vulva morphology is provided (Mok et al., 2015).

### *mlt-11* is an essential gene required for embryogenesis and molting

MLT-11 is predicted to be a large protein (234-341 kDA) with a signal sequence, a thyroglobulin domain, and 10 Kunitz protease inhibitor domains (Fig. 3, 4A). A key feature of Kunitz domains is the presence of 6 conserved cysteine residues which form three disulfide bonds critical for stabilizing the domain (Ranasinghe and McManus, 2013; Fig. 3). While Kunitz domain 8 may not be active as it is missing cysteines in the second and fourth position, the remaining Kunitz domains appear functional as they contain key conserved residues (Fig. 3). To gain insight into MLT-11 structure and function, we generated a deletion series to determine which domains were necessary for *mlt-11* function. Homozygous deletion of the signal sequence, Kunitz domains 2-10, 3-10, or 7-10 caused embryonic lethality. We balanced the mutations genetically by crossing to a strain with a *myo-2p::GFP::unc-54 3’UTR* cassette inserted into F46B3.7, a gene roughly 40kb away. We never observed progeny from balanced mutant worms lacking GFP. There was no evidence of haploinsufficiency as we could maintain balanced deletion strains. Additionally, these balanced worms produced roughly 25% dead embryos, a rate expected for a homozygous lethal mutation (Fig. 4C). In contrast, Kunitz domain 10 appeared dispensable for development as deletion animals were viable (Fig. 4C).

**Fig. 3.**
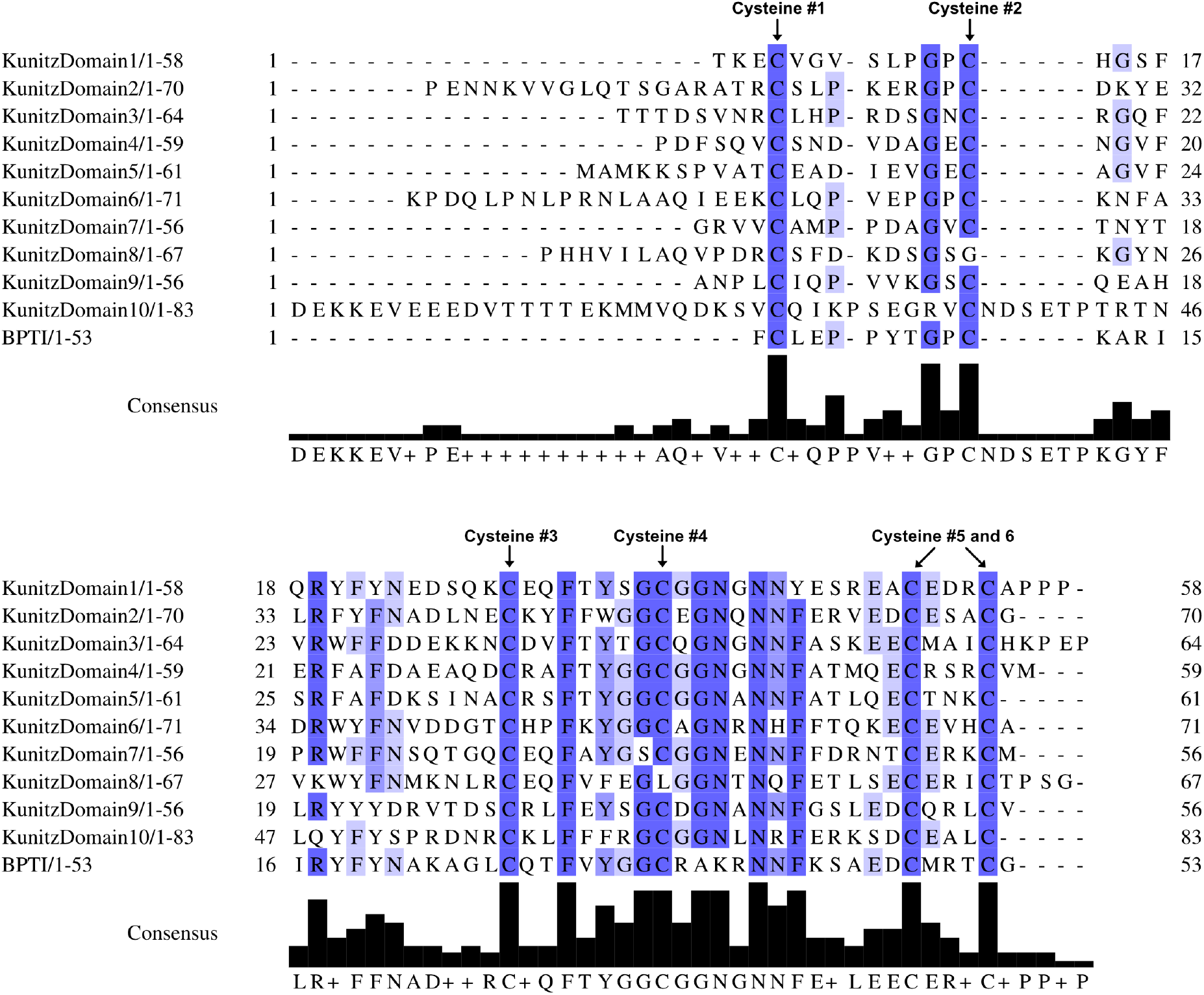
MLT-11 Kunitz domain alignment. Alignment of the ten MLT-11 Kunitz domains to Bovine Pancreatic Trypsin Inhibitor (BPTI). Positions of the six cysteine residues critical for the structure of each Kunitz domain are indicated above.

**Fig. 4.**
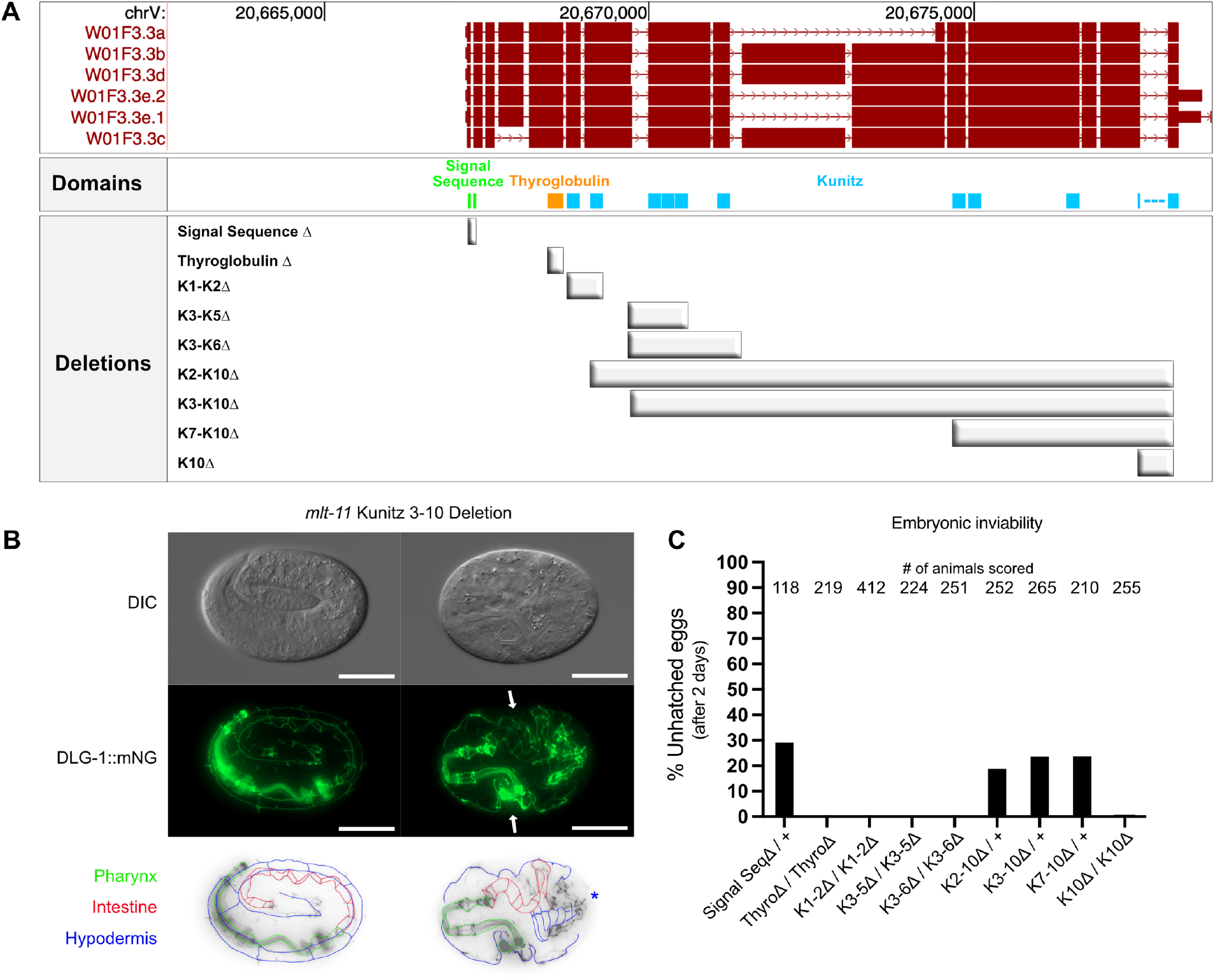
The MLT-11 C-terminus is essential for embryonic development. (A) Schematic of the *mlt-11* genomic locus with location of the indicated domains and deletions generated by genome editing. (B) Representative DIC and DLG-1::mNG images of embryos laid by a *mlt-11(*Δ*Kunitz 3-10)/+* animal. One-quarter of the embryos show the disrupted DLG-1 pattern on the right. Images are representative of 40 animals examined over two biological replicates. White arrows show invaginations of the outer membrane in homozygous mutant embryos. Blue star indicates the suspected aggregation of epithelial cells in homozygous mutant embryos. Scale bars are 20 µm. (C) Embryonic viability of the indicated *mlt-11* deletion mutants.

Deletion of Kunitz domains 1-2, 3-5 and 3-6 were completely viable, producing no dead eggs as homozygotes, but instead had coordination defects rolling to the right during forward movement. Given MLT-11::mNG expression in embryos (Fig. 2), we next examined the nature of the embryonic lethality in *mlt-11* deletion mutants using a DLG-1::mNG allele to mark adherens junctions (Heppert et al., 2018). In control embryos, DLG-1::mNG labeled adherens junctions in the pharynx, intestine, and hypodermis (Fig 4B). In contrast, Kunitz 3-10Δembryos matched to the same stage displayed severe disorganization (Fig. 4B). The pharynx and foregut adherens junctions appeared wild type, but the remainder of the junctions were disorganized and there was evidence of invaginations in the hypodermis (Fig. 4B). These data implicate the signal sequence and Kunitz 7-10 region as being essential for embryonic development.

### *mlt-11* knockdown causes defective cuticle structure and function

As we were interested in the role of *mlt-11* in promoting molting, we turned to RNAi. Both *mlt-11* (RNAi) and *mNeonGreen* (RNAi) reduced levels of MLT-11::mNG (Fig. 5A,C), and resulting phenotypes included ecdysis defects where animals were trapped in the old cuticle or failed to shed the old cuticle producing a corset (Fig. 5B). We also observed disorganized and discontinuous alae (Fig. 5B). *mlt-11* (RNAi) animals developed more slowly than control animals (Fig. 5D) and appeared to move more slowly. To test whether *mlt-11* (RNAi) caused a locomotion defect, we performed a thrashing assay, scoring body bends/minute. *mlt-11* RNAi caused nearly a 2.5-fold decrease in thrashing compared to control animals (Fig. 5E). To determine in which tissue(s) *mlt-11* was necessary to promote molting we used a set of tissue-specific RNAi strains (Fig. 6A). *mlt-11* knockdown in JDW371, a tissue-specific RNAi strain that restricts knockdown to seam, hypodermal, and intestinal cells (Johnson et al., 2022), phenocopied *mlt-11* (RNAi) in wildtype or *mlt-11::mNG* (JDW391) animals with respect to developmental delay and molting defects (Fig. 6A-C). *mlt-11* (RNAi) in QK52, a hypodermal and seam cell-specific RNAi strain, produced less penetrant developmental delay and molting defects (Fig. 6A-C). Notably, *mlt-11* (RNAi) in a hypodermal-specific RNAi strain, JDW510, produced no developmental delay or molting defects, suggesting that *mlt-11* activity is necessary in seam cells.

**Fig. 5.**
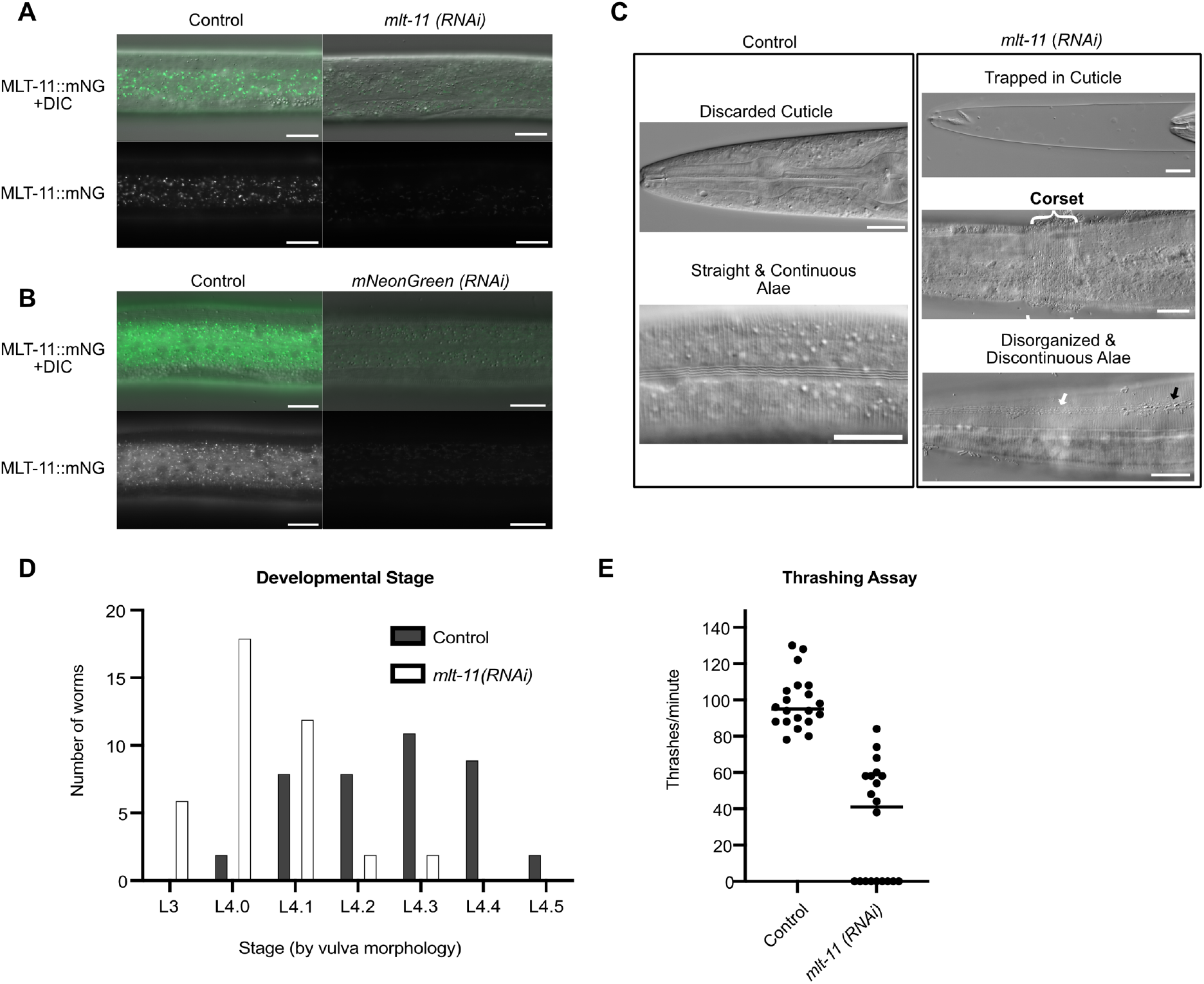
*mlt-11* knockdown causes developmental delay, molting defects, and reduction of motility. (A and B) Images of MLT-11::mNG in control, *mlt-11 (RNAi)*, or *mNeonGreen*(RNAi) animals with and without a DIC merge. (C) Representative images of molting defects in *mlt-11 (RNAi)* animals. The bracket highlights a corset phenotype, the white arrow a discontinuous alae and the black arrow, residual cuticle material stuck onto the alae following ecdysis. In A-C Scale bars are 20 µm and images are representative of 40 animals examined over two biological replicates. (D) Animals were synchronized by a timed egg lay on control or *mlt-11 (RNAi)* plates and scored for developmental stage 48 hours later using vulval development (Mok et al., 2015). Data are pooled from two independent replicates (E) 20 L4 animals grown on control or *mlt-11 (RNAi)* plates were picked into a drop of M9+gelatin and thrashes were counted for one minute.

**Fig. 6.**
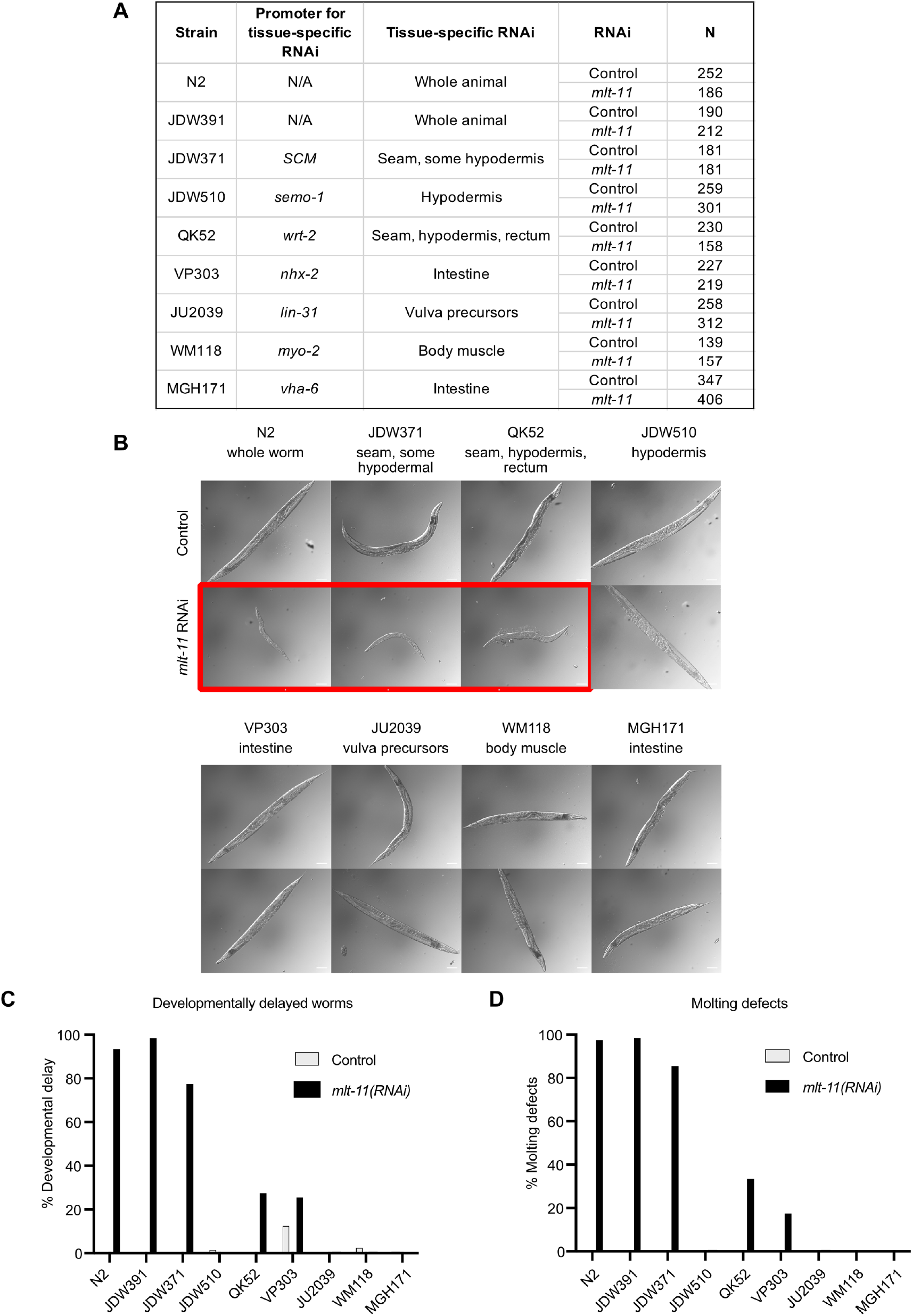
*mlt-11* is necessary in seam cells for molting and larval development. (A) Tissue-specific RNAi strains used. A timed egg lay of animals of the indicated genotype were performed on control or *mlt-11 (RNAi)* plates and phenotypes were scored three days later. (B) Representative images of control and *mlt-11 (RNAi)* on the indicated strains. The red box highlights conditions that produced smaller larvae with molting defects. (C) Developmental delay was scored in the indicated strains grown on control or *mlt-11 (RNAi)* plates and classified as a failure to reach adulthood after 72 hours of growth. (D) Molting defects were scored in the indicated strains on control or *mlt-11 (RNAi)* plates. Scored defects included animals dragging cuticles, ecdysis failure and cuticle corsets. Tissue-specific RNAi data is from two independent replicates.

### *mlt-11* inactivation causes defects in aECM structure and function

The developmental delay and molting defects caused by seam cell specific *mlt-11 (RNAi)* were reminiscent of our recent work on *nhr-23* (Johnson et al., 2022). As NHR-23 depletion causes a defect in the cuticle barrier, we tested whether *mlt-11* inactivation also compromises this barrier. We incubated control and *mlt-11* (RNAi) animals with the cuticle impermeable, cell membrane permeable Hoechst 33258 dye and scored animals with stained nuclei. In control animals we observed no Hoechst staining while in *mlt-11* (RNAi) animals 93% of animals had Hoechst-stained nuclei (Fig. 7A,B). Similar to NHR-23-depleted animals (Johnson et al., 2022), *mlt-11* (RNAi) also caused sensitivity to hypo-osmotic shock (Fig. 7C) and activation of an *nlp-29::GFP* promoter reporter activated by infection, acute stress, and physical damage to the cuticle (Fig 7D; Pujol et al., 2008; Zugasti and Ewbank, 2009).

**Fig. 7.**
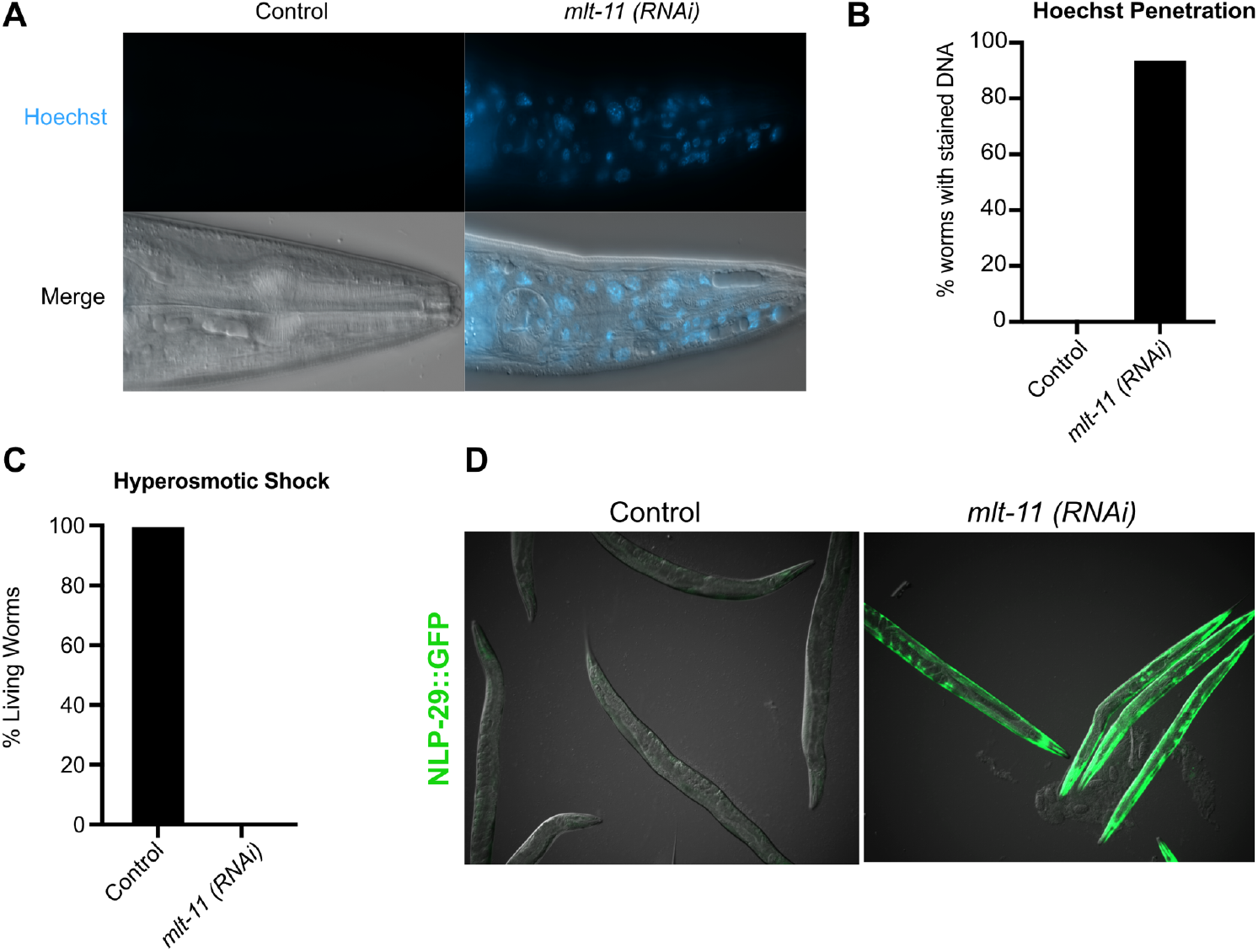
*mlt-11* knockdown causes defective aECM structure and function. (A,B)Synchronized animals were grown on control or *mlt-11 (RNAi)* plates for 48 hours before being washed off and incubated with the cuticle impermeable/membrane permeable Hoechst 33258 dye. Representative images are shown in A. Quantitation of Hoechst 33258 nuclear staining in control and *mlt-11 (RNAi)* animals. Data from 40 worms are pooled from two independent replicates. (C) Hypo-osmotic shock assay. 40 synchronized control and *mlt-11 (RNAi)* young adult animals were picked into 200 µl of dH20 and viability was scored 20 minutes later. Two biological replicates were performed and data pooled. (D) Representative images of animals carrying an *nlp-29p::GFP* reporter grown on control or *mlt-11 (RNAi)* plates. The reporter is activated by infection, acute stress, and physical damage to the cuticle (Pujol et al., 2008; Zugasti and Ewbank, 2009). Several biological replicates were performed and over 100 animals scored.

### Weak *nekl* alleles suppress *mlt-11 (RNAi)* phenotypes

We observed MLT-11::mNG localization to lysosomes and the aECM in the cuticle, rectal epithelium, vulva, and excretory duct (Fig. 2) and *mlt-11* inactivation caused defects in the aECM barrier function and localization of select aECM components (Fig. 7). MLT-11 could be acting directly in the aECM or could function in lysosomes, as this organelle has been shown to play an important role in aECM remodeling during molting (Miao et al., 2020). To distinguish between these possibilities, we examined the genetic interaction between *mlt-11* (RNAi) and weak *nekl* alleles. *nekl-2* and *nekl-3* encode NIMA-related kinases that regulate endocytosis and are required for completion of molting (Joseph et al., 2020; Laźetić and Fay, 2017a; Yochem et al., 2015). Weak *nekl-2* and *nekl-3* hypomorphs are viable but display reduced clathrin-mediated endocytosis (Joseph et al., 2020). We reasoned that if MLT-11 acted in the aECM then weak *nekl* alleles might suppress the *mlt-11* (RNAi) defects by trapping more MLT-11 in the aECM. Conversely, if MLT-11 activity was necessary in lysosomes then weak *nekl* alleles might enhance the *mlt-11* (RNAi) defects by reducing the amount of MLT-11 that is trafficked to lysosomes. Weak *nekl-2(fd81)* and *nekl-3(gk894345)* alleles suppressed the small body size of *mlt-11* (RNAi) animals (Fig. 8A-C) and suppressed *mlt-11* (RNAi) molting defects (Fig 8D).

**Fig. 8.**
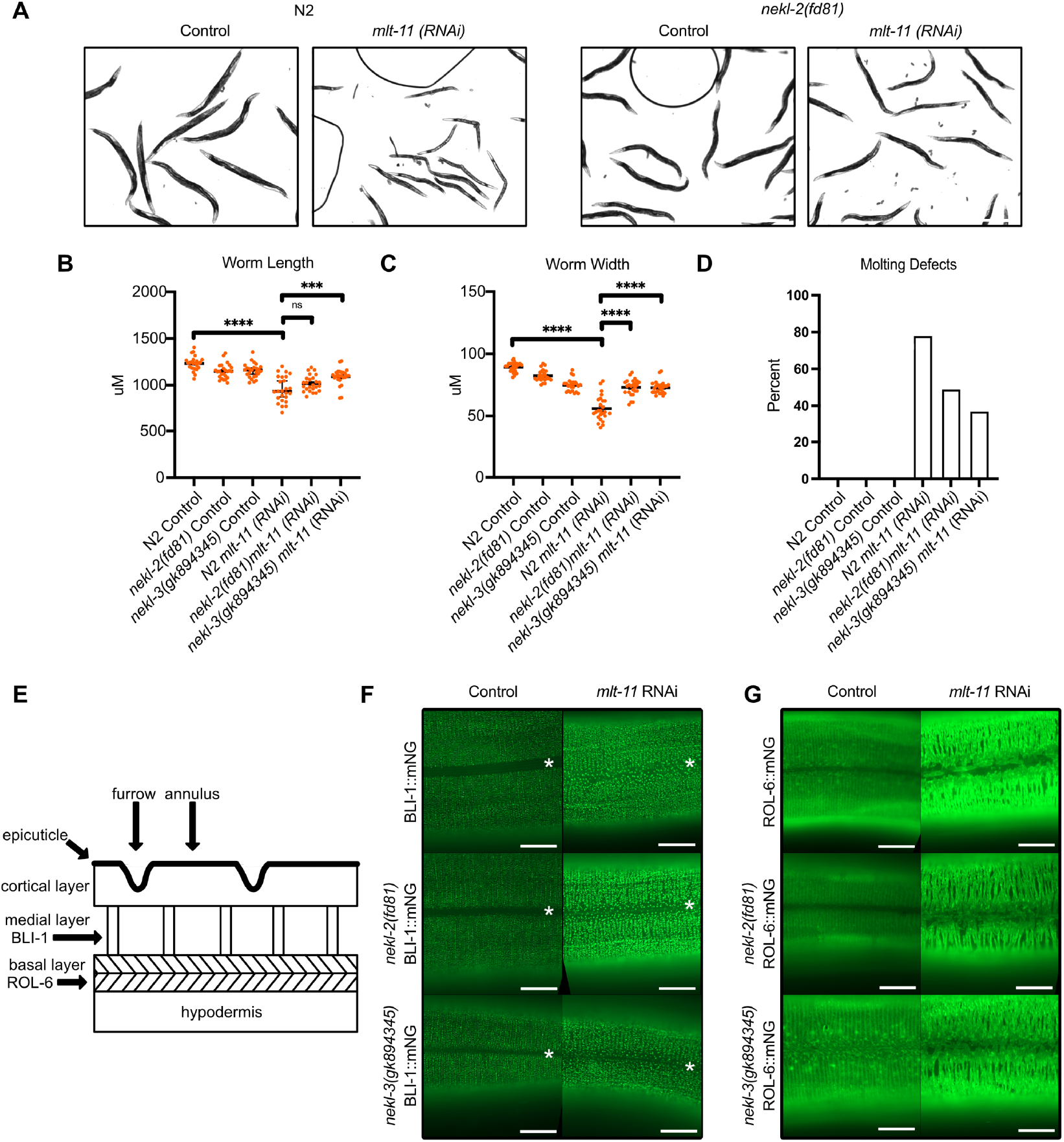
Weak *nekl* alleles suppress *mlt-11 (RNAi)* defects. (A) Representative images of animals of the indicated genotypes grown on control or *mlt-11 (RNAi)* plates. Average length (B) and width (C) of animals of the indicated genotypes grown on control or *mlt-11 (RNAi)* plates. Horizontal black lines indicate average from 2 independent experiments. Synchronized animals of the indicated genotypes were grown on control or *mlt-11 (RNAi)* plates and scored for the number that displayed molting defects (D). Schematic displaying the primary layers and structures making up the *C. elegans* cuticle including the localization of ROL-6 and BLI-1 collagens used in (F) and (G). Representative images of *rol-6::mNG* (F) and *bli-1::mNG* (G) animals grown on control or *mlt-11 (RNAi)* plates. Scale bars are 20 µm. Two independent experiments were performed and a minimum of 30 worms observed. In F, white stars indicate where the seam cells and alae are located.

NHR-23 depletion causes reduced levels and mis-localization of the medial cuticle layer strut collagen BLI-1 and defective localization of the basal layer collagen ROL-6 (Johnson et al., 2022), so we examined the effect of *mlt-11* (RNAi) on these markers. In control animals, BLI-1::mNG localized to regularly spaced punctae in rows and was excluded from the area of the aECM over seam cells (Fig. 8F). *mlt-11* (RNAi) caused BLI-1 to localize to larger, irregularly spaced punctae which were also found over the seam cells (Fig. 8F). This exclusion zone and BLI-1::mNG organization pattern were partially restored in weak nekl allele worms grown on *mlt-11* (RNAi). ROL-6::mNG, in control animals, localized to striped annuli, with an irregular but tight zipper-like pattern over seam cells (Fig. 8G). *mlt-11* (RNAi) animals displayed thick and aggregated ROL-6::mNG with a large gap over seam cells where left and right side extensions typically meet (Fig. 8G). Weak nekl allele worms treated with *mlt-11* (RNAi) had a similar aggregation of ROL-6::mNG over hypodermal cells, but more frequent connections across seam cells to ROL-6::mNG in annuli on the opposite side. These data indicated that *mlt-11* is necessary for aECM structure and MLT-11 acts in the aECM to promote development and molting.

## DISCUSSION

How aECMs are dynamically remodeled during development and disease remains poorly understood. Using the *C. elegans* cuticle as a model aECM we demonstrate a role for the protease inhibitor MLT-11 in promoting embryogenesis, molting, wild type developmental rate, and the aECM barrier. MLT-11::mNG oscillates and localizes to the aECM in the larval cuticle, vulva, rectum, and excretory pore, and is also in lysosomes. In embryos, MLT-11::mNG is secreted into the space between the eggshell and the embryo and then localizes to the cuticle prior to hatching. Tissue-specific RNAi data indicates that *mlt-11* primarily acts in seam cells. Depletion of *mlt-11* results in mislocalization of the collagens ROL-6 and BLI-1, and genetic data suggests that MLT-11 functions primarily in the aECM.

We observed three distinct phenotypes depending upon the severity of *mlt-11* mutation or depletion: i) embryonic lethality; ii) larval molting defects and developmental delay; and iii) rolling. Any C-terminal deletions removing Kunitz domains 7-10, including Kunitz 2-10 and Kunitz 3-10 deletions, produced embryonic lethality (Fig. 4). DLG-1::mNG revealed severe disorganization of adherens junctions in these mutants with defects being most pronounced in the hindgut and hypodermis (Fig. 4). One possibility is that MLT-11 is required for embryonic sheath function. The embryonic sheath is an aECM that preserves embryonic integrity and distributes force during embryo elongation (Kelley et al., 2015; Vuong-Brender et al., 2017). MLT-11 is secreted during the window of morphogenesis, when the embryo elongates. One model is that MLT-11 restrains protease activity to ensure sheath integrity during elongation and in its absence the sheath is compromised. Inactivation of sheath components has been shown to cause embryo arrest and rupturing (Vuong-Brender et al., 2017). It is unclear whether the molting defects and developmental delay incurred by *mlt-11* (RNAi) reflect a distinct molecular defect or arise from a similar role for MLT-11 during larval aECM remodeling. The collagens ROL-6 and BLI-1 exhibited aberrant localization in *mlt-11* (RNAi) treated larvae. A conditional deletion approach would be ideal to create a *mlt-11* null in larvae, bypassing the embryo phenotypes.

Our Kunitz 1-2, 3-5 and 3-6 deletions all produced a weak right roller phenotype. The mapping locus *rol-9* was recently discovered to be encoded by a gain-of-function *mlt-11* allele (Rich et al., 2022). How does a protease inhibitor mutation cause a roller phenotype? Aside from *mlt-11*, the only non-collagen roller gene is *rol-3*, which encodes a predicted receptor tyrosine kinase (Jones et al., 2013). ROL-3 is hypodermally expressed and necessary for ecdysis and cuticle formation (Jones et al., 2013). *rol-3* mutations cause defects in seam cell formation and *mlt-11* is necessary in seam cells for developmental progression and molting (Fig. 6). One possibility is that weak *mlt-11* alleles provide sufficient activity to promote ecdysis but elevated protease activity disrupts collagen processing, leading to a roller phenotype. Interestingly, *mlt-11* (RNAi) disrupts ROL-6::mNG localization and specific alleles of both *mlt-11* and *rol-6* cause a right roller phenotype. In the future it will be interesting to test whether *mlt-11* and *rol-3* genetically interact and whether *mlt-11* inactivation affects the localization of collagens that when mutated produce left roller phenotypes.

Transcription factors regulate complex networks of genes to control cellular and developmental processes. Assigning the regulation of a single regulated gene to a phenotype incurred by inactivation of a given transcription is challenging. NHR-23 depletion causes developmental delay, molting defects, and defective aECM structure and barrier function (Johnson et al., 2022). Strikingly, *mlt-11* (RNAi) phenocopies NHR-23 depletion in many regards. Both cause developmental delays, apolysis defects, and a loss of the aECM barrier function (Fig. 5-7; Johnson et al., 2022). The ROL-6::mNG localization defects are highly similar, with annular disorganization and a gap over the seam cells (Fig. 8; Johnson et al., 2022). Tissue-specific RNAi indicates that the seam cells are a key site of action for both *nhr-23* and *mlt-11*, though *nhr-23* activity also appears necessary in hypodermal cells (Fig. 6; Johnson et al., 2022). NHR-23-regulated genes are enriched in protease inhibitors (Johnson et al., 2022), and *mlt-11* is a critical gene for promoting aECM remodeling during molting (Fig. 8). An open question is whether MLT-11 is unique in mediating the NHR-23-dependent molting program or whether these terminal phenotypes are a common feature of disrupting components in the NHR-23 gene regulatory network. Given that *mlt-11* is a protease inhibitor gene, the common phenotypes suggest that some aspects of the NHR-23 depletion phenotype may be due to unrestrained protease activity. Identifying which protease(s) that MLT-11 inhibits and the protease substrates is a critical future direction.

Why does MLT-11 have so many Kunitz domains? The extensively studied bovine pancreatic trypsin inhibitor has a single Kunitz domain (Ascenzi et al., 2003), as do other proteins such as Alzheimer Precursor Protein (Beckmann et al., 2016). Others such as Tissue Factor Pathway Inhibitor and *C. elegans* MEC-9 have multiple Kunitz domains (Broze and Girard, 2012; Du et al., 1996). One possibility was that the large number of Kunitz domains in *C. elegans* MLT-11 arose through recent duplication. Arguing against this possibility many *Caenorhabdid* species, as well as more distantly related nematodes (*P. pacificus, O. vovlulus, B. malayi*) have large MLT-11 homologs with predicted signal sequences and 10 Kunitz domains (Fig. S1). In *C. elegans*, Kunitz domains 1-6 and 10 appear dispensable whereas deletion of Kunitz domains 7-9 causes embryonic lethality (Fig. 4). Notably, there is additional sequence flanking Kunitz domain 9 that is conserved (Fig. S1). Interestingly, our immunoblotting experiments detect a smaller isoform of 50-70 kDa that could be produced by cleavage at or near the start of Kunitz domain 9. An interesting approach would be to exogenously express a C-terminal fragment of *mlt-11* containing Kunitz domains 7-10 in worms with an endogenous null allele of *mlt-11* to see if this region is sufficient to rescue embryonic inviability. We would reasonably expect these rescued worms to be right rollers as our deletion strains lacking Kunitz 1-2, 3-5 and 3-6 are right rollers.

Our data could suggest that the different Kunitz domains may play distinct roles, or that their location within the protein is important. Kunitz domains work as competitive protease inhibitors, which would suggest that MLT-11 could serve as a scaffold to bind to and inactivate proteases. Kunitz domains tend to inactivate serine proteases, yet there are no serine proteases implicated in molting (Frand et al., 2005). *nas-37*, an astacin metalloprotease, peaks in expression 30 minutes after *mlt-11* mRNA peaks in expression and genes expressed at similar points in development often function in common processes (Davis et al., 2004; Farrell et al., 2018; Hendriks et al., 2014; Meeuse et al., 2020). MLT-11 may regulate uncharacterized protease inhibitors or could inactivate different classes of protease inhibitors. An unusual family of Kunitz domain protease inhibitors from the parasitic nematode *Fasciola hepatica* was shown to inhibit cathepsin proteases, not serine proteases (Smith et al., 2020). Alternatively, MLT-11 may not function as a protease inhibitor. The Kunitz domain containing molting factor BLI-5 was shown to enhance the activity of two serine proteases, rather than inhibit them (Stepek et al., 2010). Similarly, the ADM-2 protease regulates molting by modulating levels of the low-density lipoprotein receptor–related protein, LRP-1, through a mechanism independent of its protease activity (Joseph et al., 2022).

### Future perspective

Our characterization of MLT-11 provides an entry point into understanding how proteases and protease inhibitors interact to promote aECM remodeling. Going forward, exploring whether MLT-11 plays roles in specialized aECM such as the vulval lumen, excretory duct, and glial socket cuticle will be important. As proteases are important targets to combat parasitic nematode infections, understanding how they are regulated during development by endogenous protease inhibitors will be critical to develop novel approaches to combat this group of devastating pathogens.

## MATERIALS AND METHODS

### Strains and culture

*C. elegans* were cultured as originally described (Brenner, 1974), except worms were grown on MYOB media instead of NGM. MYOB agar was made as previously described (Church et al., 1995).

### Strains created by injection in the Ward Lab and used in this study

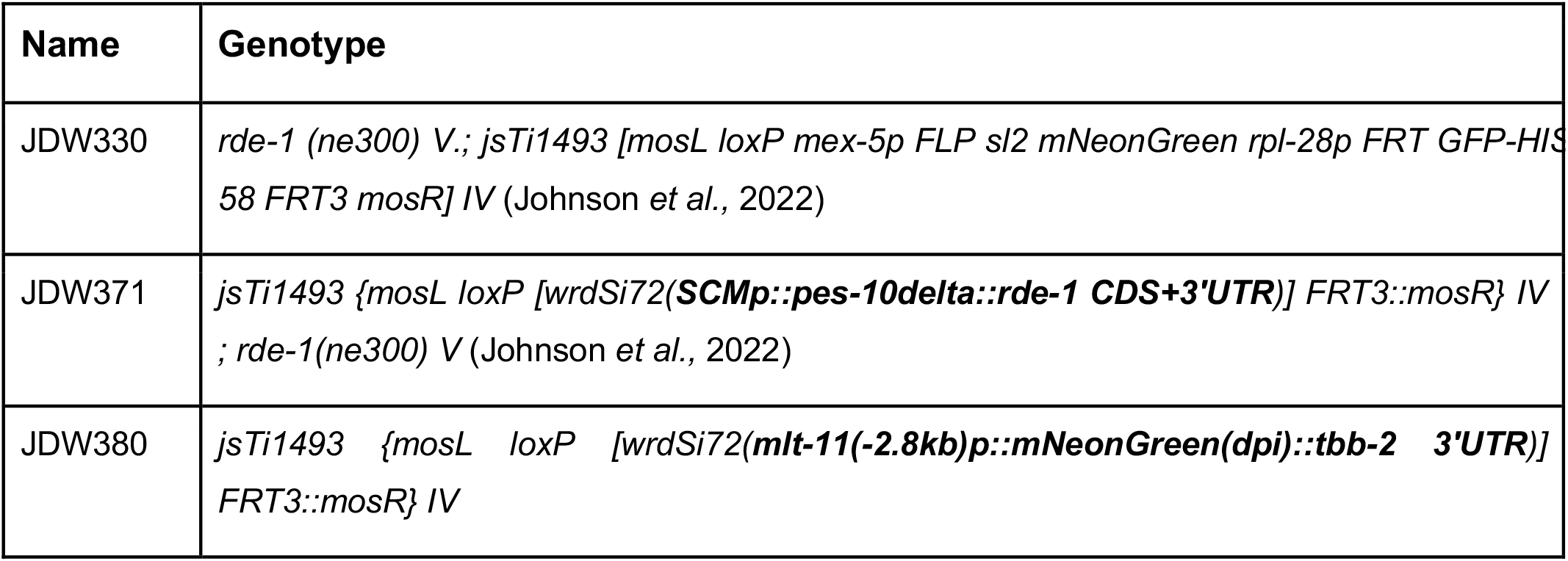
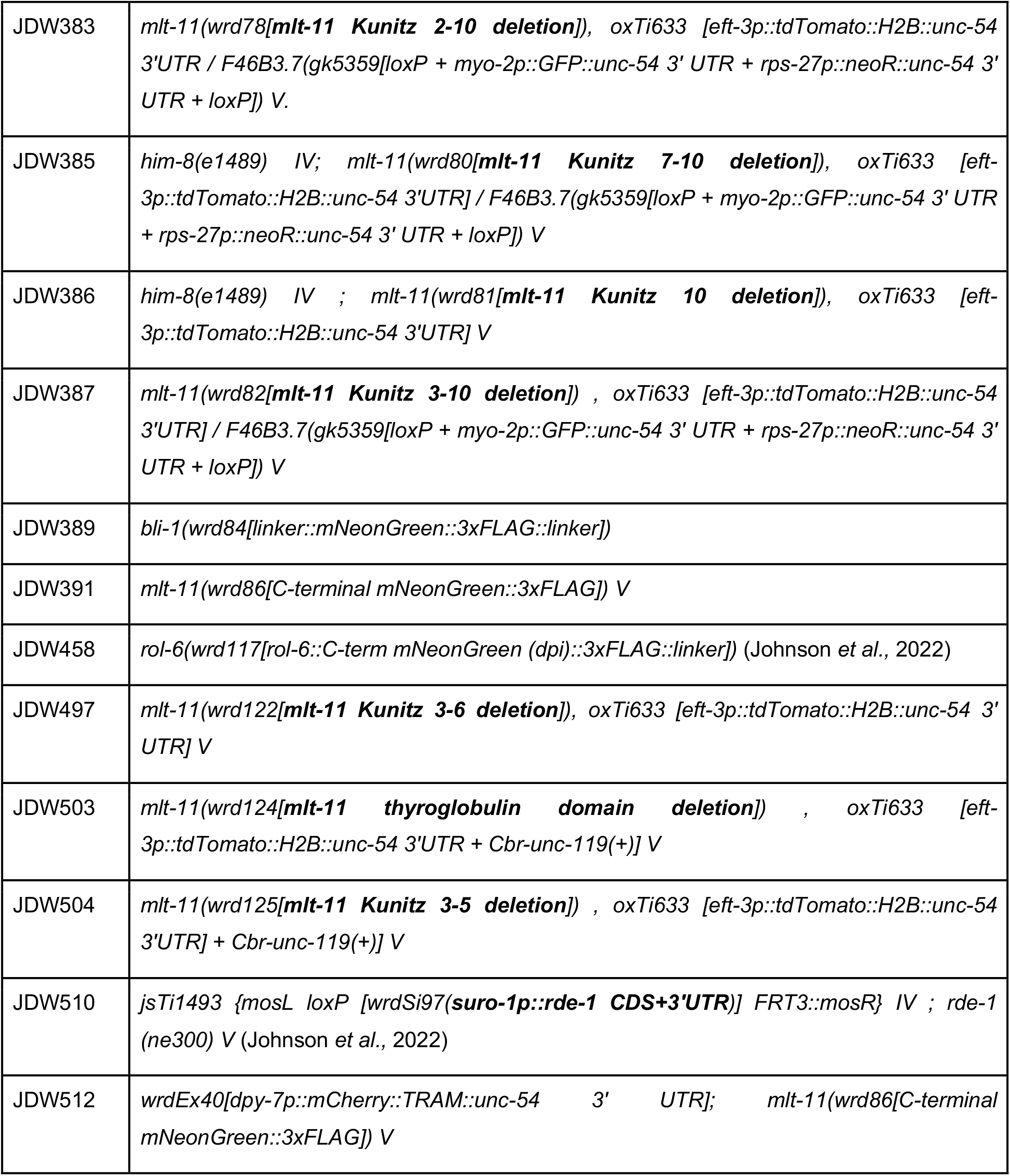

### Strains created by crossing in the Ward Lab and used in this study

**Strains created by crossing in the Ward Lab and used in this study:**
| Strain | Genotype | Crossing strain 1 | Crossing strain 2 |
| --- | --- | --- | --- |
| JDW359 | *ezIs2 III [fkh-6::GFP + unc-119(+)] ; him-8(e1489) IV ; F46B3.7(gk5359[loxP + myo-2p::GFP::unc-54 3' UTR + rps-27p::neoR::unc-54 3' UTR + loxP]) V.* | DZ325 | VC4276 |
| JDW401 | *mlt-11(wrd82[mlt-11 Kunitz 3-10 deletion]) , oxTi633 [eft-3p::tdTomato::H2B::unc-54 3'UTR] / F46B3.7(gk5359[loxP + myo-2p::GFP::unc-54 3' UTR + rps-27p::neoR::unc-54 3' UTR + loxP]) V. dlg-1(cp301[dlg-1::mNG-C1^3xFlag]) X* | JDW387 | LP598 |
| JDW511 | *ezIs2 [fkh-6::GFP + unc-119(+)] / + III ; him-8(e1489) / + IV ; mlt-11(wrd126[mlt-11 signal sequence deletion...mNG::3xFLAG]) / F46B3.7(gk5359[loxP + myo-2p::GFP::unc-54 3' UTR + rps-27p::neoR::unc-54 3' UTR + loxP]) V* | EG7968 | JDW359 |
| JDW516 | *bli-1(wrd84[linker::mNeonGreen::3xFLAG::linker]) II ; nekl-3(gk894345) X* | WY1141 | JDW389 |
| JDW517 | *rol-6(wrd117[rol-6::C-term-mNeonGreen (dpi)::3xFLAG::linker]) II ; nekl-3(gk894345) X* | WY1141 | JDW458 |
| JDW522 | *nekl-2(fd81); rol-6(wrd117[rol-6::C-term-mNeonGreen (dpi)::3xFLAG::linker]) II* | WY1122 | JDW458 |
| JDW526 | *nekl-2(fd81);bli-1(wrd84[linker::mNeonGreen::3xFLAG::linker]) II* | WY1122 | JDW389 |

### Strains provided by the *Caenorhabditis* Genetics Center

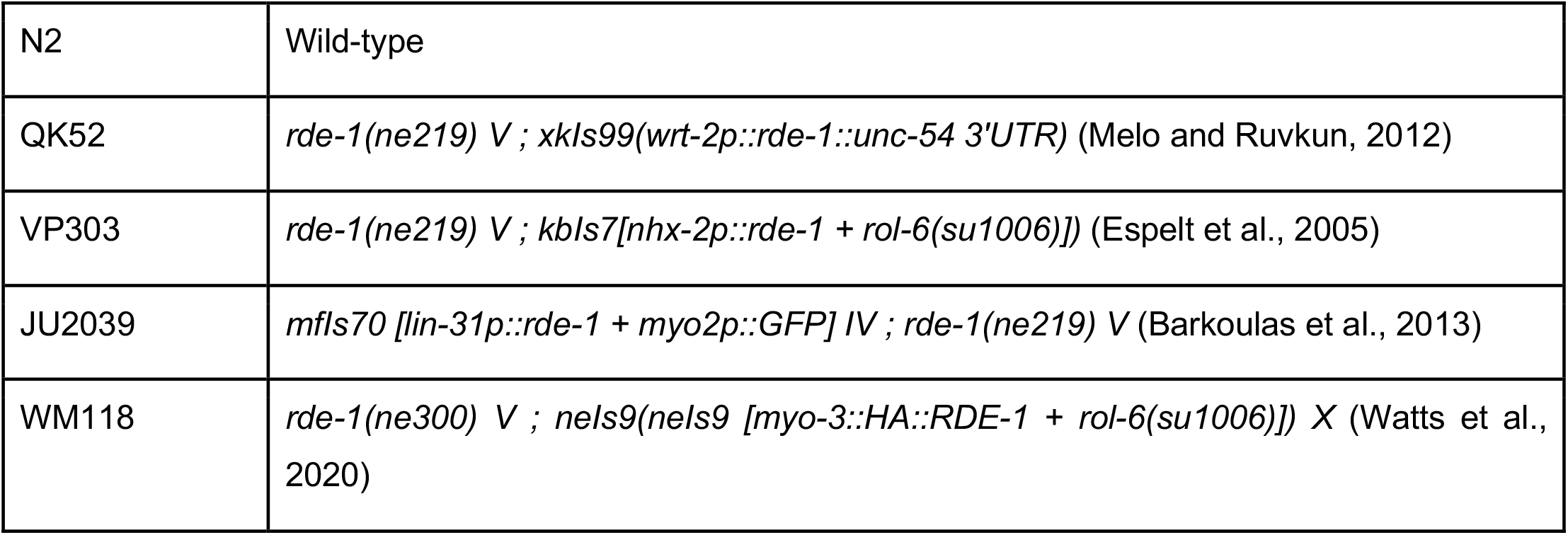
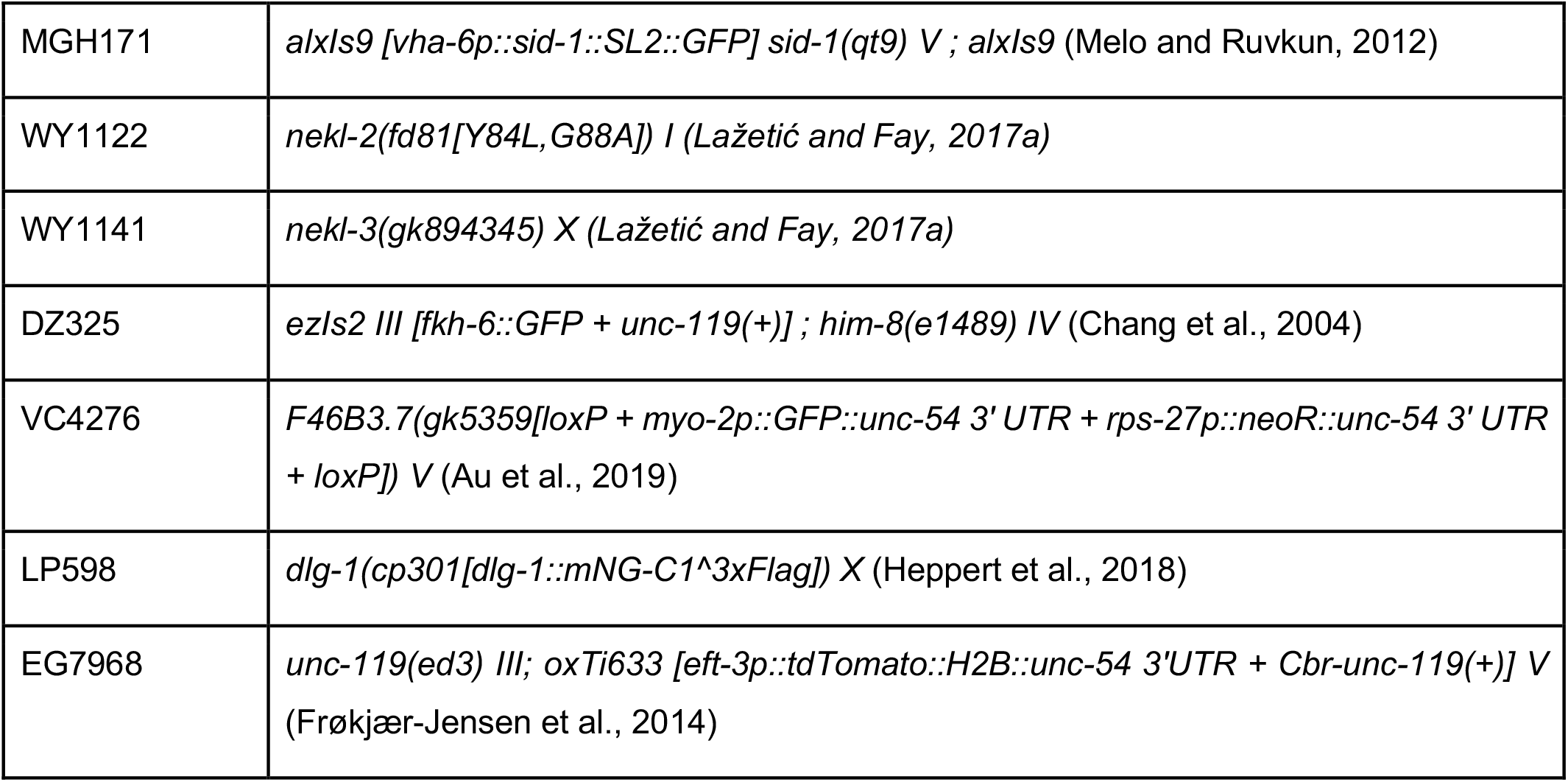

### Other strains

NM5548 *jsSi1579 jsSi1706 jsSi1726[loxP myo-2p FRT nlsCyOFP myo-2 3’ mex-5p FLP D5 glh-2 3’ FRT3] II* was a gift from Dr. Michael Nonet and will be described elsewhere. The sequence of this landing pad can be found on the Nonet lab website (https://sites.wustl.edu/nonetlab/rmce-insertion-strains-lastedited6-4-2022/) and is inserted at an sgRNA within 50 base pairs away from the ttTi5605 insertion site.

### Genome Editing

All plasmids used are listed in Table S1. Annotated plasmid sequence files are provided in File S1. Specific cloning details and primers used are available upon request. JDW380 *jsTi1493 {mosL loxP [wrdSi72(****mlt-11(−2*.*8kb)p::mNeonGreen(dpi)::tbb-2 3’UTR****)] FRT3::mosR} IV* was created by recombination-mediated cassette exchange (RMCE)(Nonet, 2020). A 2.8 kb *mlt-11* promoter fragment was initially Gibson cloned into the *NLS::mScarlet (dpi)::tbb-2 3’UTR* vector pJW1841 (Ashley et al., 2021) to generate pJW1934. The mScarlet cassette was then replaced with mNeonGreen (dpi) to generate pJW2229. The *mlt-11p (−2*.*8kb) mNeonGreen (dpi)-tbb-2 3’UTR* fragment was PCR amplified from pJW2229 and Gibson cloned into *Sph*I-HF+*Spe*I-HF double digested RMCE integration vector pLF3FShC to produce pJW2337. This vector was integrated into NM5179 and the SEC was excised as previously described (Nonet, 2020).

*mlt-11* deletion strains were created by injection of Cas9 ribonucleoprotein complexes (RNPs)(Paix et al., 2014; Paix et al., 2015) [700 ng/µl IDT Cas9, 115 ng/µl each crRNA and 250 ng/µl IDT tracrRNA], oligonucleotide repair template (110 ng/µl) and pSEM229 co-injection marker (25 ng/µl)(El Mouridi et al., 2020) for screening into strain EG7968. Where possible, we selected “GGNGG” crRNA targets as these have been the most robust in our hand and support efficient editing (Farboud and Meyer, 2015). F1s expressing the co-injection marker were isolated to lay eggs and screened by PCR for the deletion. F2 progeny of a verified F1 deletion mutant were crossed to JDW359 males expressing *myo-2::GFP* to genetically balance the mutation. Genotyping primers are provided in Table S2. JDW389 and JDW391 and were created by injection of RNPs [700 ng/µl IDT Cas9, 115 ng/µl crRNA and 250 ng/µl IDT tracrRNA] and a dsDNA repair template (25-50 ng/ul) created by PCR amplification of a plasmid template into N2 animals (Paix et al., 2014; Paix et al., 2015)(Table S1). PCR products were melted to boost editing efficiency, as previously described (Ghanta and Mello, 2020). For the *mlt-11* C-terminal knock-in, the mNeonGreen::3xFLAG cassette was inserted right at the double-strand break and a stop codon followed the 3xFLAG sequence. We re-coded the sequence between the insert and native stop codon and placed it in 5’ to the mNeonGreen 3xFLAG insertion (File S2). Sequences of CRISPR/Cas9-mediated genome edits are provided in File S2. crRNAs used are provided in Table S3. F1 progeny were screened by mNeonGreen expression. JDW512 *wrdEx40[dpy-7p::mCherry::TRAM::unc-54 3’ UTR]; mlt-11(wrd86[C-terminal mNeonGreen::3xFLAG]) V* was generated by injection of a *dpy-7p::mCherry::tram-1::unc-54 3’UTR* vector (25 ng/µl)(Chen et al., 2012) into JDW391. F1 progeny were screened by mCherry::TRAM expression.

### Imaging

Synchronized animals were collected from MYOB, control, or auxin plates by either picking or washing off plates. For washing, 1000 µl of M9 + 2% gelatin was added to the plate or well, agitated to suspend animals in M9+gelatin, and then transferred to a 1.5 ml tube. Animals were spun at 700xg for 1 min. The media was then aspirated off and animals were resuspended in 500µl M9 + 2% gelatin with 5 mM levamisole. 12 µl of animals in M9 +gel with levamisole solution were placed on slides with a 2% agarose pad and secured with a coverslip. For picking, animals were transferred to a 10 µl drop of M9+5 mM levamisole on a 2% agarose pad on a slide and secured with a coverslip. Images were acquired using a Plan-Apochromat 40x/1.3 Oil DIC lens or a Plan-Apochromat 63x/1.4 Oil DIC lens on an AxioImager M2 microscope (Carl Zeiss Microscopy, LLC) equipped with a Colibri 7 LED light source and an Axiocam 506 mono camera. Acquired images were processed through Fiji software (version: 2.0.0-rc-69/1.52p). For direct comparisons within a figure, we set the exposure conditions to avoid pixel saturation of the brightest sample and kept equivalent exposure for imaging of the other samples.

### Western Blot

For the western blot in Fig. X JDW391 animals were synchronized by alkaline bleaching (dx.doi.org/10.17504/protocols.io.j8nlkkyxdl5r/v1) and released on MYOB plates. Animals were harvested at the indicated time points by picking thirty animals into 30 µl of M9+0.05% gelatin. Laemmli sample buffer was added to 1X and then samples were immediately incubated for five minutes at 95°C. Lysates were stored at -80°C until resolution by SDS-PAGE. Lysates were resolved using precast 4-20% MiniProtean TGX Stain Free Gels (Bio-Rad) with a Spectra™ Multicolor Broad Range Protein Ladder (Thermo; # 26623) protein standard. For the anti-FLAG blots, proteins were transferred to a polyvinylidene difluoride membrane by wet transfer using Towbin buffer (25 mM Tris, 192 mM glycine, 20% methanol, pH 8.3) supplemented with 0.1% SDS and 30V was applied for 16 hours in a cold room. The buffer was chilled prior to use and a freezer back was added to the transfer container. For the anti-Tubulin blots, a semi-dry transfer with a TransBlot Turbo (Bio-Rad) was performed. Blots and washes were performed as previously described (Johnson et al., 2022, 23). Anti-FLAG blots used horseradish peroxidase (HRP) conjugated anti-FLAG M2 (Sigma-Aldrich, A8592-5×1MG, Lot #SLCB9703) at a 1:2000 dilution. Mouse anti-alpha-Tubulin 12G10 (Developmental Studies Hybridoma Bank; “-c” concentrated supernatant) was used at 1:4000 and Digital anti-mouse (Kindle Biosciences LLC, R1005) diluted 1:20,000 was used as the 2°. Blots were incubated for 5 minutes with 1 ml of Supersignal West Femto Maximum Sensitivity Substrate (Thermo Fisher Scientific, 34095) and the final blot were imaged using the ‘chemi high-resolution’ setting on a Bio-Rad ChemiDoc MP System.

### RNAi Knockdown

RNA interference experiments were performed as in Johnson *et al*. (2022). Control RNAi used either an empty L4440 or high-efficiency T444T RNAi vector (Sturm et al., 2018). The *mlt-11 (RNAi)* vector was streaked from the Ahringer library (Kamath et al., 2003). The *mNeonGreen*(RNAi) vector was generated by synthesizing a cDNA fragment and cloning it into T444T. Synthesis and cloning were performed by Twist Bioscience. Vector sequences are provided in File S1.

## Acknowledgements

We would like to thank Doug Kellogg and Tyler DeWitt for helpful conversations. We thank Zoe Johnson, Javier Hernandez Lopez, Zoie Reyna, Emma Cadena, Valarie Hallin, and Olivia Vedar for research support. Some strains were provided by the Caenorhabditis Genetics Center, which is funded by the NIH Office of Research Infrastructure Programs [P40 OD010440]. The anti-alpha tubulin 12G10 monoclonal antibody developed by J. Frankel and E.M. Nelson of the University of Iowa was obtained from the Developmental Studies Hybridoma Bank, created by the NICHD of the NIH and maintained at The University of Iowa, Department of Biology, Iowa City, IA 52242.

## Author Contributions

Conceptualization: J.M.R, J.D.W.

Methodology: J.M.R, J.D.W.

Validation: J.M.R, J.D.W.

Formal analysis: J.M.R, J.D.W.

Resources: J.M.R, J.D.W.

Data curation: J.M.R, J.D.W.

Writing - original draft: J.M.R, J.D.W.

Writing - review & editing: J.M.R, J.D.W.

Supervision: J.M.R, J.D.W.

Project administration: J.D.W.

Funding acquisition: J.D.W.

---

**Figure S1.**
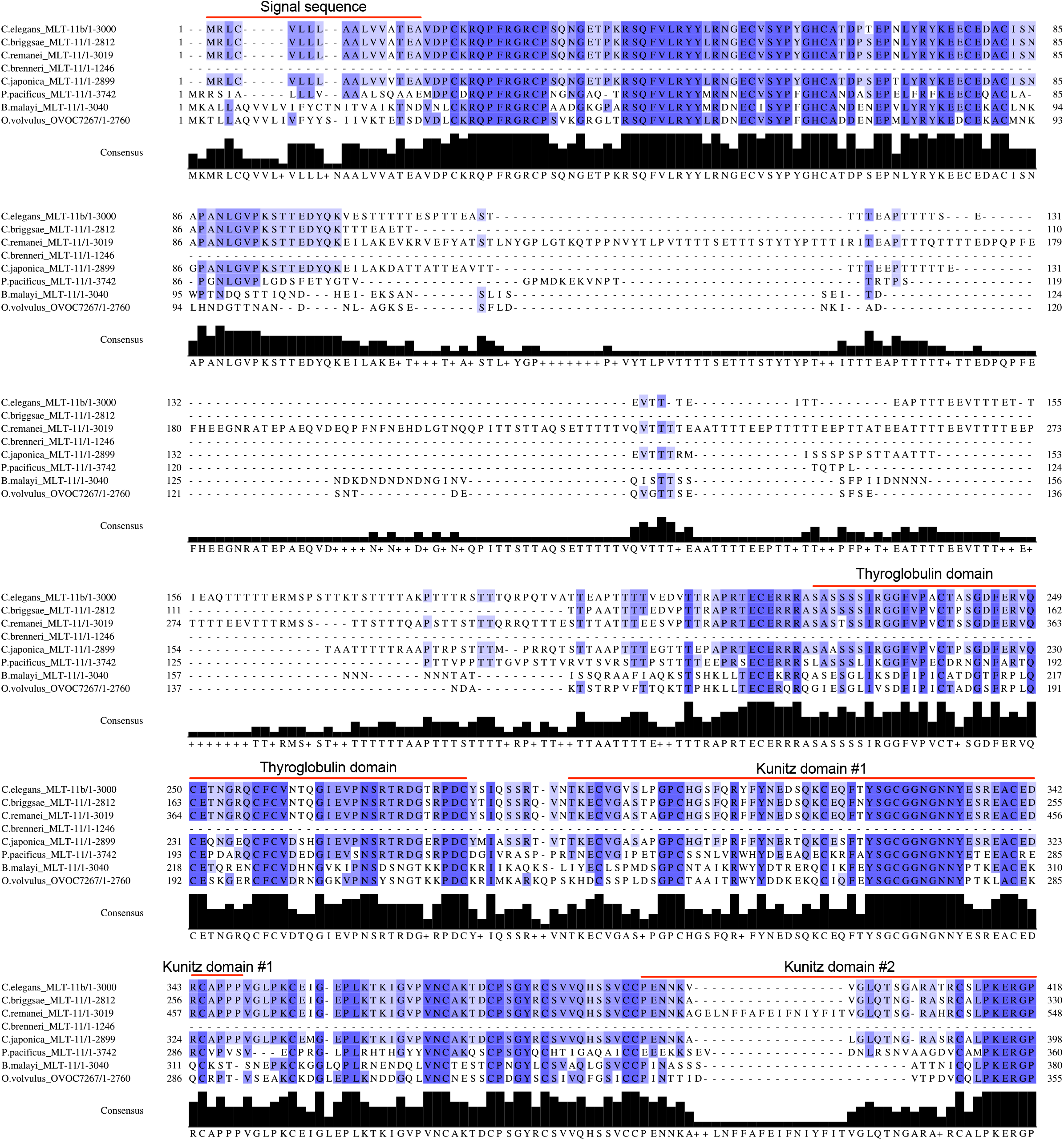
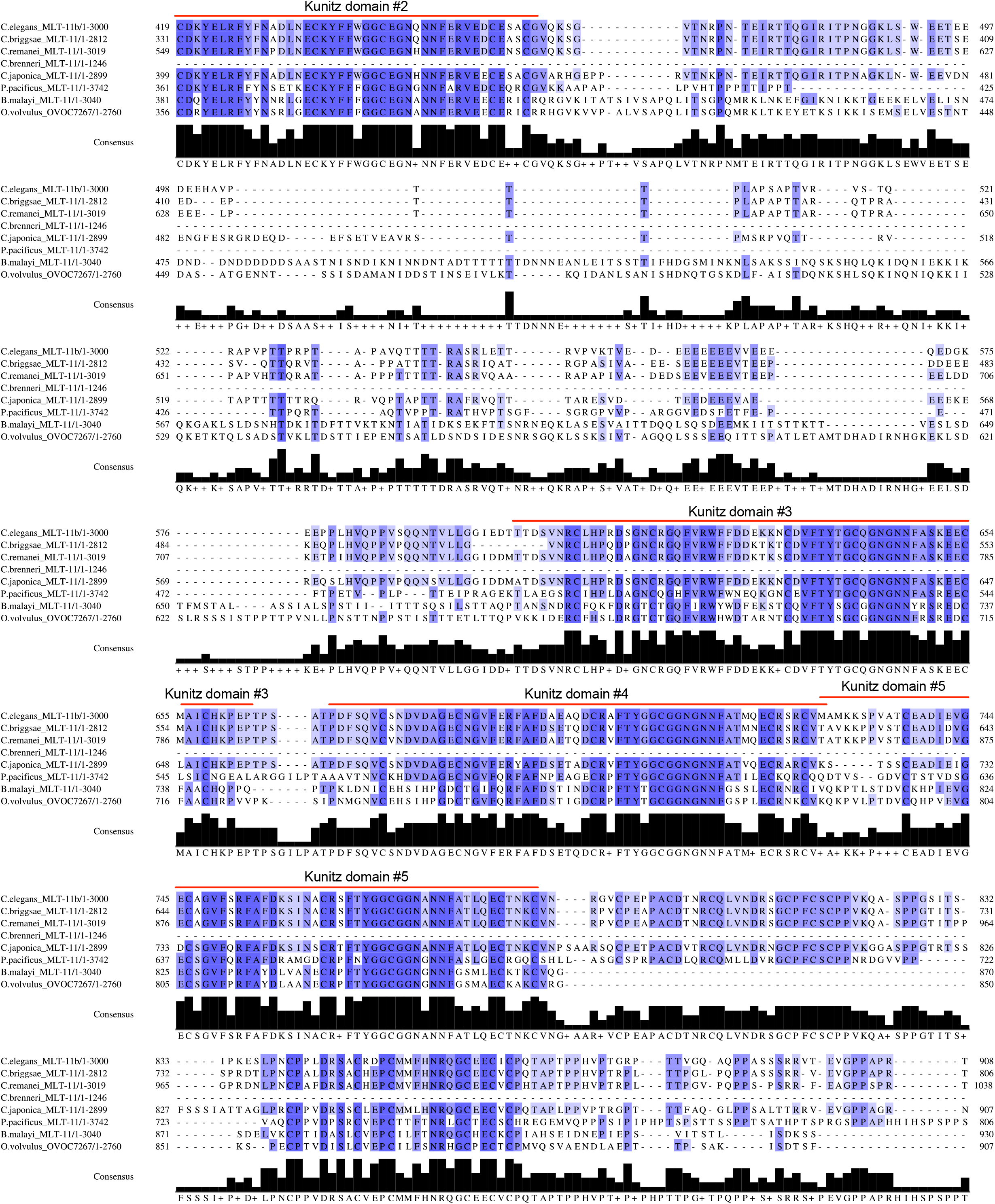
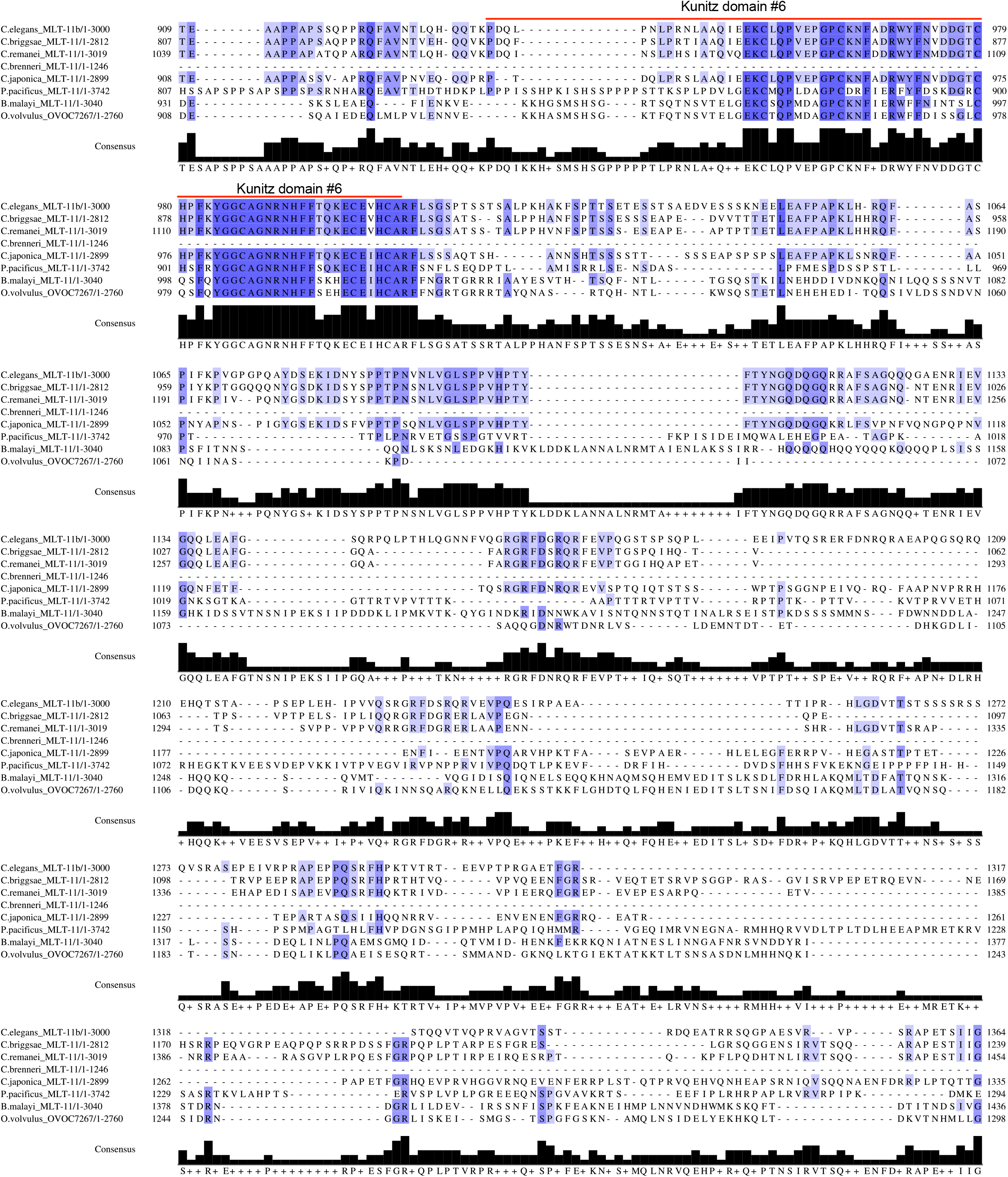
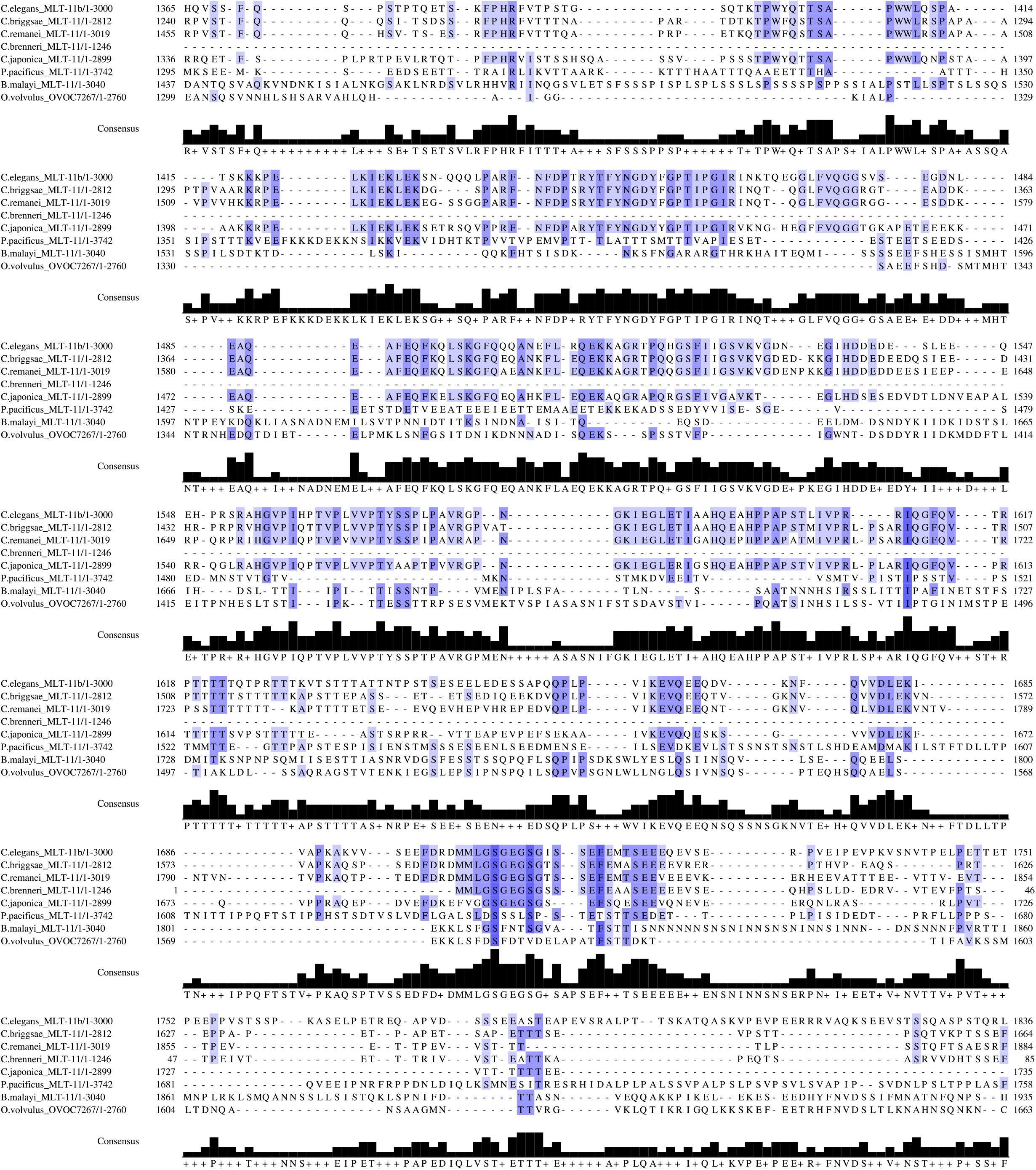
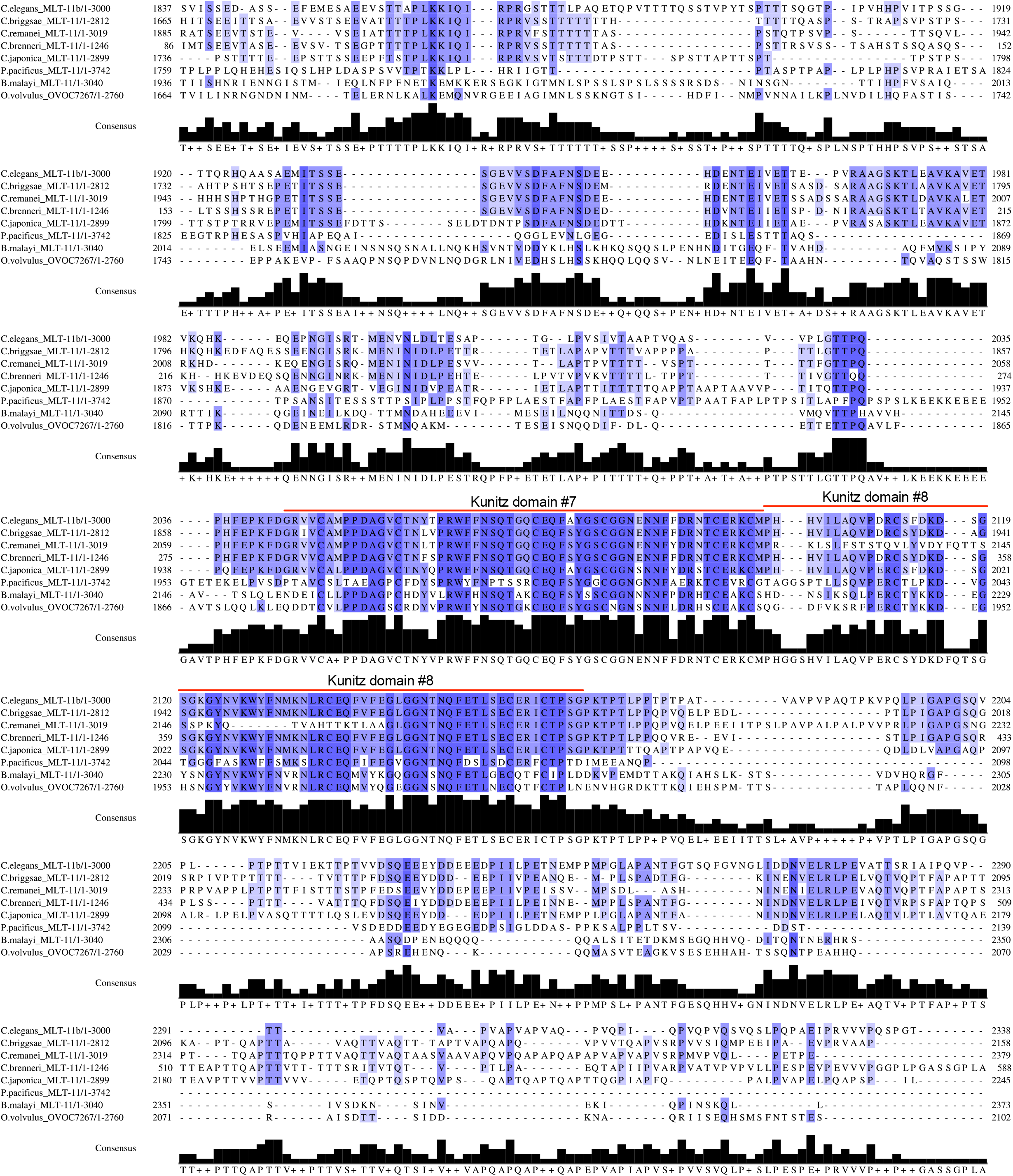
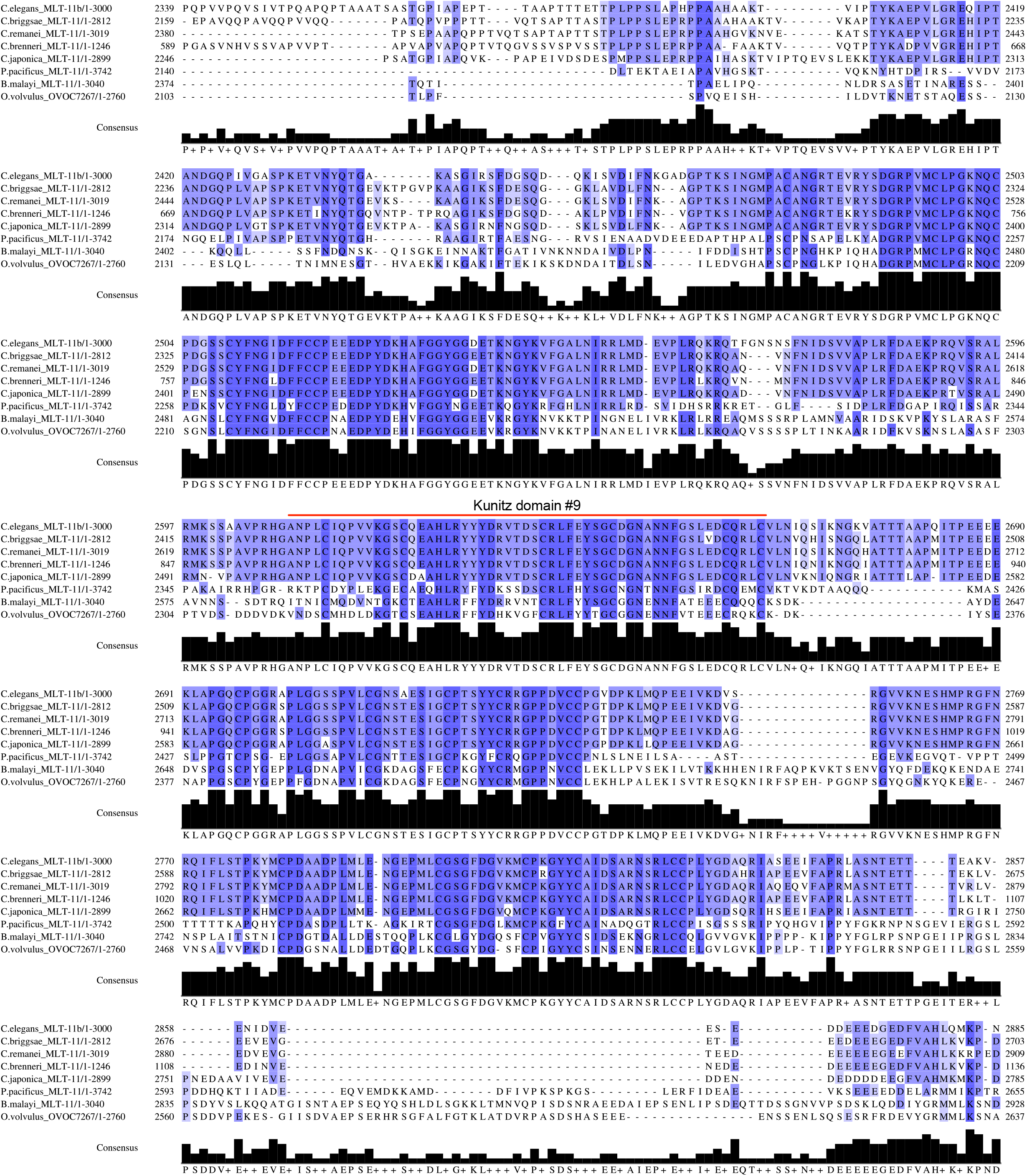
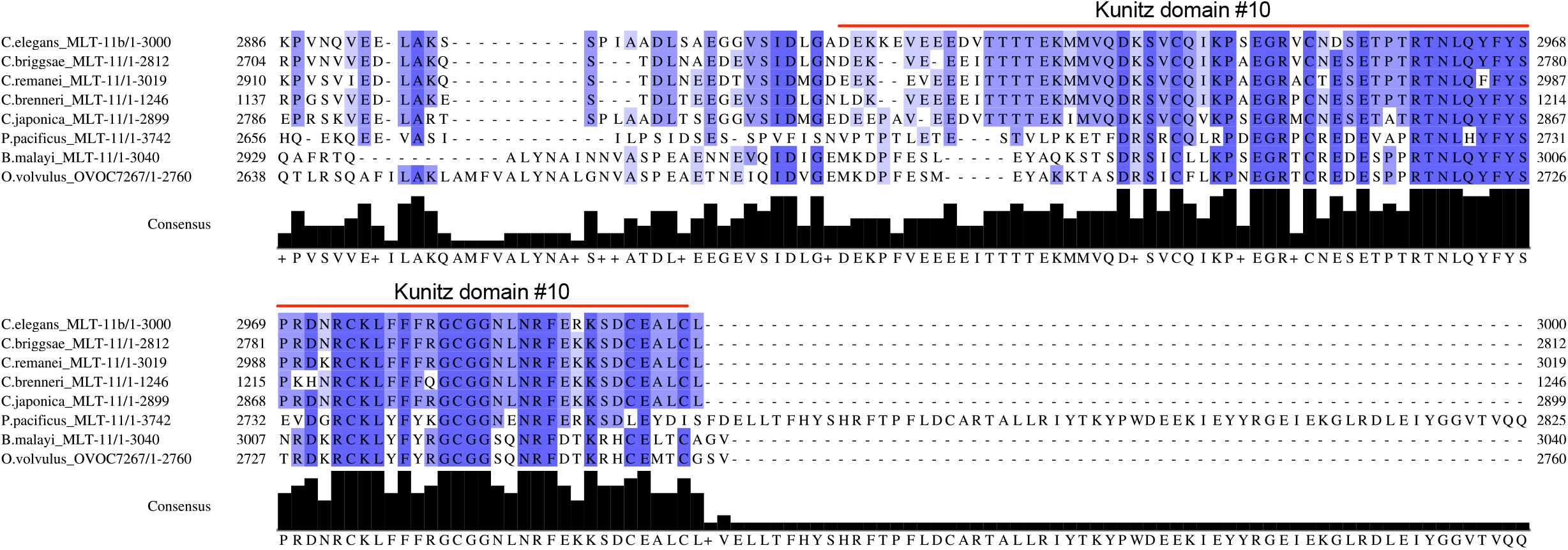
Alignment of MLT-11 homologs. MLT-11 homologs from the indicated nematode species were aligned using Clustal Omega. The length in amino acids of each homolog follows the species and homolog name. To the left and right of the alignment are amino acid positions of the end residues for each protein. Blue shading indicates conserved sequences and the histogram at the bottom depicts the degree of conservation with a consensus sequence listed below. The positions of the *C. elegans* MLT-11 signal sequence, thyroglobulin domain, and ten Kunitz domains are indicated. We note that we cut off the extended *P. pacificus* C-terminus (amino acids 2626-3742) since no sequence aligned to it as all proteins terminated at the *C. elegans* MLT-11 stop codon. No predicted motifs are found in the *P. pacificus* C-terminus.

